# A deep-learning predictive model for Cas12a activity in wheat

**DOI:** 10.64898/2026.11.18.767513

**Authors:** Christophe Gaillochet, Pieter Wytynck, Alexandra Pena Fernandez, Ellen Slaman, Jonas De Saeger, Myriam Shafie, Debbie Rombaut, Els Van Lerberge, Tingting Zhu, Ward Develtere, Lieven Sterck, Marnik Vuylsteke, Denys Marushchak, Ive De Smet, Dirk Inzé, Laurens Pauwels, David De Vleesschauwer, Frank Meulewaeter, Thomas B. Jacobs

## Abstract

CRISPR technology has created targeted DNA mutations across the tree of life. Target sites are defined by RNA molecules that largely determine the efficiency of an experiment. Several predictive models are available to select the most efficient RNAs. Unfortunately, these models are not built with plant data and have limited predictive power in plant systems. To overcome this limitation, we established a high-throughput arrayed wheat protoplast transfection method and measured the editing efficiency of ∼1,000 unique LbCas12a crRNAs at on- and off-target sites. We trained different models by testing 40 deep learning architectures and found that a convolutional neural network model called 2CaP-5 displayed the best performance. While 2CaP-5 had moderate to strong predictive power in protoplasts, it did not generalize to *Agrobacterium*-mediated transformation. Despite this limitation, we demonstrate that 2CaP-5 can be re-trained with as few as 500 crRNAs. The use of such a small training set is advantageous for low-throughput experimental systems such as typically found in transgenic plant systems. We also used ∼2,000 off-targets in the wheat subgenomes to investigate Cas12a specificity. Our dataset confirms previous observations regarding off-targets, specifically that single mismatches reduce indel efficiency provided they are adjacent to the PAM and that off-target activity can be largely prevented by three mismatches. Altogether, our results show that the 2CaP-5 model has moderate to strong predictive power for Cas12a crRNAs in wheat protoplasts and will serve as a useful foundation for predictive tools in other plant species.

## Background

CRISPR-mediated mutagenesis has become a routine method to introduce precise mutations in genomes. The CRISPR-associated (Cas) enzymes Cas9 and Cas12a (previously Cpf1) are widely used in the gene editing field. Both enzymes are two-component systems, consisting of the Cas endonuclease and a guide RNA (gRNA; Cas9) or CRISPR RNA (crRNA; Cas12a) that directs the Cas protein to its genomic target. One difference between these Cas endonucleases is the protospacer adjacent motif (PAM) needed for target recognition. While SpCas9 recognizes 5’-NGG-3’ PAM targets and introduces double-stranded breaks (DSBs) that are either blunt or possess a 1-nt 5’ overhang, LbCas12a targets a 5’-TTTV-3’ PAM and generates DSBs with 4-5 nt 5’ overhangs leading to larger deletions than Cas9 [1]. Despite the relatively fewer target sites available for Cas12a than for Cas9, Cas12a has garnered particular interest among the agriculture biotechnology sector given its clearer IP landscape [2].

A key determinant for successful mutagenesis is gRNA/crRNA efficiency. High-throughput studies have shown that different gRNAs can result in widely different mutation frequencies and DNA repair outcomes [3]. In plants, where the generation of mutant lines takes months to years and has limited output, it is highly desirable to accurately predict nuclease activity to increase the chances of experimental success. CRISPR screens would particularly benefit from more efficient guide selection as even modest gains in efficiency can drastically decrease the size of the screening population [4].

Several predictive models have been developed for gRNA efficiency in animal systems and these have been instrumental for increasing efficiency. For Cas9, several well-known tools are the Azimuth score [5], FORECast [3], inDELPHI [6], and DeepSpCas9 [7, 8]. For Cas12a, CINDEL [9], DeepCpf1 [10], CRISPR-DT [11] and a Random Forest model [12] have been reported. These models were trained with data from large screens in animal systems typically containing thousands of datapoints. While such predictive models are accurate in their respective systems, they often do not generalize to different organisms or experimental methods [13]. In plants, a low correlation was found for 22 Cas9 target sites in tobacco using eight different predictive tools [14]. A screen of 89 gRNAs in tomato found that the Azimuth score could predict the top and bottom performers to some extent, but in general the correlation was weak [15] and there were similar observations with DeepCpf1 [16]. These studies show that, despite the extensive development of CRISPR tools in plant biotechnology, there is currently no method to accurately predict gRNA efficiency in plants.

Wheat (*Triticum aestivum*) is an important staple crop. It is cultivated on more than 240 million ha worldwide and constitutes over 30% of the total human caloric intake, making it a prime candidate for crop improvement through genome editing [17]. Modern wheat varieties have a relatively large (∼14.7 Gb) and repetitive allohexaploid (AABBDD) genome [18, 19]. The three subgenomes are highly homologous, complicating CRISPR-mediated genome editing experiments. Targeting all homoeoalleles of a single gene on all three subgenomes requires efficient multiplexing, while specifically targeting a single subgenome is hampered by potential off-targeting at the non-target homoealleles. The ability to reliably predict gRNA/crRNA efficiencies and off-target editing would greatly help alleviate these design challenges.

A major difficulty in developing predictive models in plants is the lack of large-scale datasets reporting gRNA/crRNA efficiencies [20, 21]. Generating the number of transformed plants necessary to accurately determine mutation efficiencies for such large numbers would require substantial resources. Although several large, pooled knockout screens encompassing thousands of gRNAs have been performed in rice, maize and Arabidopsis, tomato and cotton [22–27], the number of regenerated plants per gRNA in such studies is too low to accurately determine gRNA efficiencies and in most instances researchers only investigate lines with phenotypes. Studies that do regenerate sufficient numbers of individuals per gRNA often only report tens of gRNAs, which is too small a dataset for predictive modeling. In contrast, protoplast systems are an attractive alternative to stable plant transformation, offering higher throughput and shorter experimental turnaround times and can even be used to generate transgene-free, edited plants in several species [28].

Here, we report the development of an arrayed CRISPR screen in wheat protoplasts to build a predictive model for LbCas12a activity. Using an initial dataset of ∼1,000 crRNAs, we assessed LbCas12a editing activity and repair products at scale. We trained and compared several deep learning models using convolutional neural networks (CNN) or recurrent neural networks (RNN) and identified one called 2CaP-5 that predicts crRNA activity with high confidence (Spearman ρ = 0.67) in wheat protoplasts using ribonucleoprotein (RNP) delivery. We further tested if the model could be generalized to plasmid transfection in protoplasts and stably transformed plants and found good correlation (Spearman ρ = 0.83) between RNP and plasmid delivery in protoplasts. However, there was a low correlation (Spearman ρ = 0.22 – 0.32) between the protoplast systems and stable transformation with *Agrobacterium*. Using publicly available LbCas12a datasets, we demonstrate that 2CaP-5 can be efficiently retrained with as few as 500-1,000 crRNAs. Thus, our results demonstrate that 2CaP-5 is a powerful predictive model for crRNA activity in wheat protoplasts and can be readily retrained on datasets of limited size, which is particularly beneficial for plant transformation.

## Results

### Target design

To develop a deep learning model for crRNA efficiency prediction in plants, we first generated a suitable dataset. We designed crRNAs to perfectly target one subgenomic site while having off-targets with mismatches in the other two subgenomes. This design not only allowed us to determine editing efficiencies at target sites, but also gain valuable information on off-target activity. We first identified 1,639 homeologous gene triplets with a single member on each subgenome. For each of these genes, we selected a random subgenome to be the on-target site, while ensuring that on-target sites were equally distributed over the different subgenomes. We selected all possible 23-nt target sites preceded by a 5’-TTTV PAM, resulting in 53,454 candidate crRNAs. From these, we selected those with 1-3 mismatches in one or both of the off-target homeologs. To simplify genotyping, we selected crRNAs where we could amplify on-and off-targets simultaneously using a single primer pair (**Fig. 1a**). Genotyping primers were tested by Illumina sequencing on wild-type samples and analyzed with CRISPResso2. Of the 1,056 primer pairs tested, 988 were considered successful since they had >500 reads for each individual homeolog and less than 2% background editing rate at the on-target sites (**Fig. S1, Additional File 1 all amplicons, Additional File 2 primers**).

**Fig. 1:**
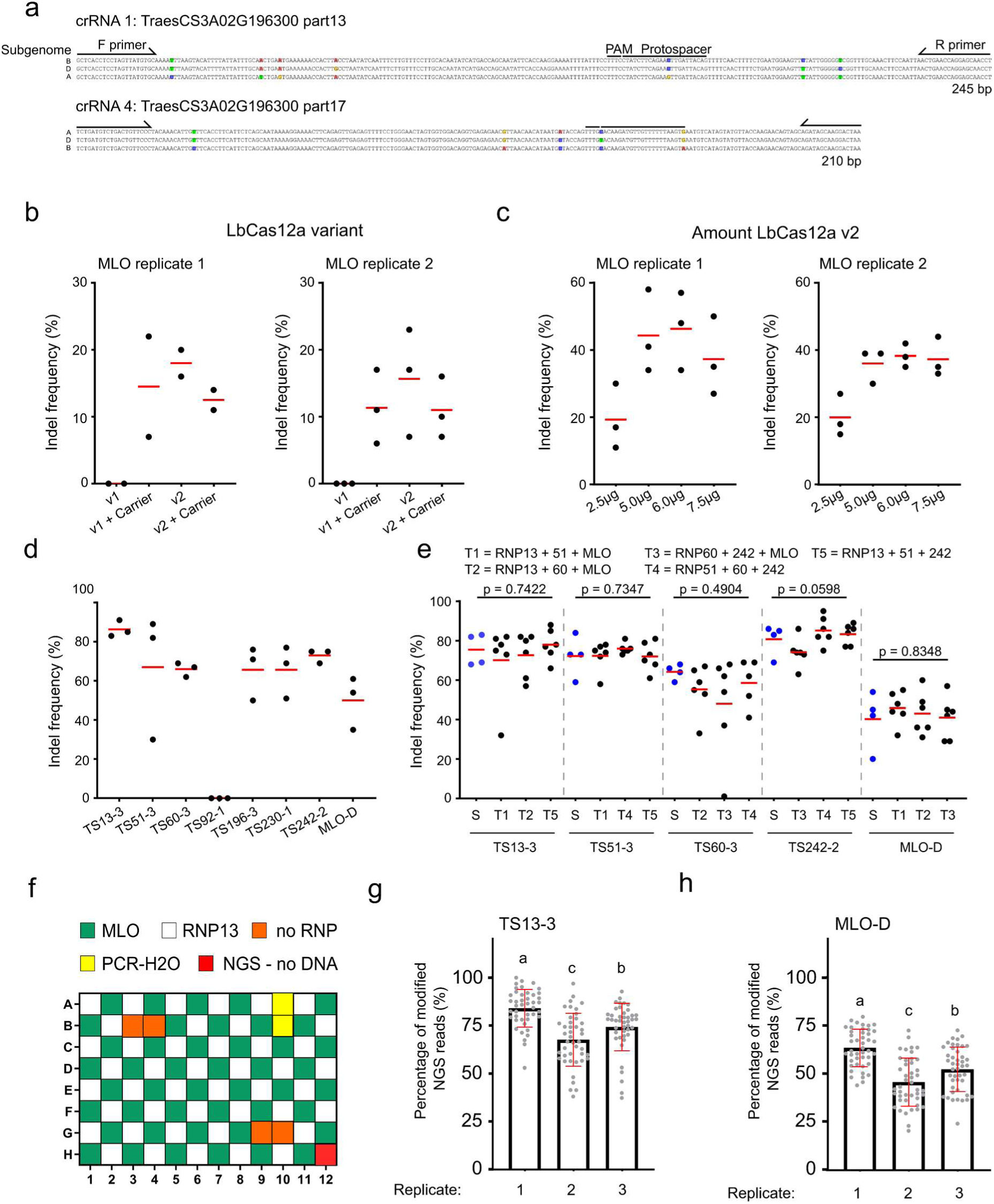
Establishment of arrayed CRISPR screen in wheat protoplasts. **(a)** Two examples of on-and off-targets. Cas12a protospacer adjacent motifs (PAM), spacer, and genotyping primers are indicated. Single nucleotide polymorphisms are indicated by highlighted bases. **(b-c)** ICE indel frequencies at TaMLO-D by different LbCas12a RNP variants (b) and protein concentrations (c). Carrier indicates the presence of a carrier plasmid. **(d)** ICE indel frequencies at 8 target sites using 6 µg of LbCas12a v2. **(e)** Comparison of simplex (one RNP per transfection - blue dots) and triplex editing (3 RNPs mixed per transfection - black dots) at five target sites. The 5 unique triplex RNP combinations (T1-T5) were used. **(f)** Schematic of the 96-well control plate transfection with RNPs targeting TS13-3 or TaMLO-D. **(g-h)** Editing efficiency at on-targets TS13-3 (g) and MLO-D (h) in independent transfection replicates of the control plate. Error bars represent standard deviation. Statistics: e,g-h: One-way ANOVA, Tukey-HSD p<0.05.

### Establishment of an arrayed CRISPR screen in wheat protoplasts

We next established a high-throughput wheat protoplast transfection protocol. We decided to transfect RNPs to eliminate the need for cloning and purifying large numbers of plasmids. We tested several parameters that might affect RNP activity including the enzyme variant and the amount of enzyme. We compared two commercially available LbCas12a variants (v1 and v2, Integrated DNA Technologies) and tested the impact of co-transfection with a carrier plasmid. We targeted the wheat *MLO* gene on subgenome D (TaMLO-D) and determined indel frequencies using Sanger sequencing and ICE analysis [29]. We did not detect indels with v1, but v2 produced average indel frequencies of 16-18% (**Fig. 1b**). While the carrier plasmid increased indel frequencies with v1, we did not observe a similar effect with v2. Based on these results, we decided to use v2 without carrier as it simplified the experimental setup. The use of 5 to 7.5 µg of protein per transfection resulted in average indel frequencies of ∼40% whereas 2.5 µg had an average efficiency of 20% (**Fig. 1c**). Thus, we decided to use 6 µg for the screen.

We targeted seven additional sites and measured average indel frequencies ranging from 50 -80% for six of them, showing that we could induce mutations at multiple loci with high efficiency and consistency (**Fig. 1d**). To increase throughput, we tested the effect of multiplexing crRNAs by comparing simplex and triplex editing for five combinations of RNPs (**Fig. 1e**). All crRNAs displayed similar indel frequencies in simplex and triplex combinations (p>0.05; One-Way ANOVA, Tukey-HSD test), indicating that multiplexing three RNPs does not affect efficiency and can be used to increase the number of RNPs tested per transfection. To test our protocol in a multi-well plate format, we performed RNP transfections in 96-well plates and measured indel efficiencies using Illumina sequencing and CRISPresso2 analysis. We alternated the transfection of two RNPs (RNP13 and MLO) across the plate and included different negative controls to assess background indel rates (**Fig. 1f**). All target sites (3 per RNP) were assessed at every position across the plate. Across all replicates, TS13-3 and MLO-D had an average background editing frequency of 0.9% and 2.2%, respectively (**Fig. S2a**). For both RNPs, we observed a significant difference in editing efficiency between the days of transfection (p<0.05, One-way ANOVA, T>2 likelihood test for dropping fixed terms), indicating that the protoplast batch/replicate is a source of variability (**Fig. 1g-h, Fig. S2b**). We also detected a significant difference between columns that varied between transfection replicates (**Fig. S2d-i**, p<0.05 One-way ANOVA, T>2 likelihood test for dropping fixed terms). Based on these results, we decided to use technical replicates and a generalized linear mixed model to account for the experimental effects.

### CRISPR screen

For the full screen, we designed twelve 96-well plates, containing 1,152 crRNAs (**Additional File 3 crRNAs**). The RNPs were transfected in triplex and each triplex was transfected in three independent protoplasts batches. We included the MLO-D crRNA as an internal control at random positions on individual plates (**Fig. S3a**). While fully randomizing the plate position of all crRNAs would have been ideal to control for row and column effects, it would have tripled the cost of crRNA synthesis and was therefore cost-prohibitive. Three target-specific PCRs, corresponding to the three targets in triplex, were performed on each sample and all amplicons were pooled and sequenced with Illumina sequencing (**Fig. S3a**). The percentages of modified reads for on- and off-targets were determined with CRISPResso2 (**Fig. 2a**). We observed consistent percentages of modified reads between each of the transfection triplicates with Pearson r = 0.91 - 0.99 between replicates (**Fig. S3b,c**). To account for batch and replicate effect, we normalized the data by fitting a Generalized Linear Mixed Model. The normalized editing efficiencies had a high correlation with the raw data (Pearson r = 0.99), indicating low variability between batches and replicates (**Fig. S3d**). During the analysis we identified 11 crRNAs that produced ambiguous reads (cannot distinguish between on- and off-target amplicons) and these were removed from later steps.

**Fig. 2:**
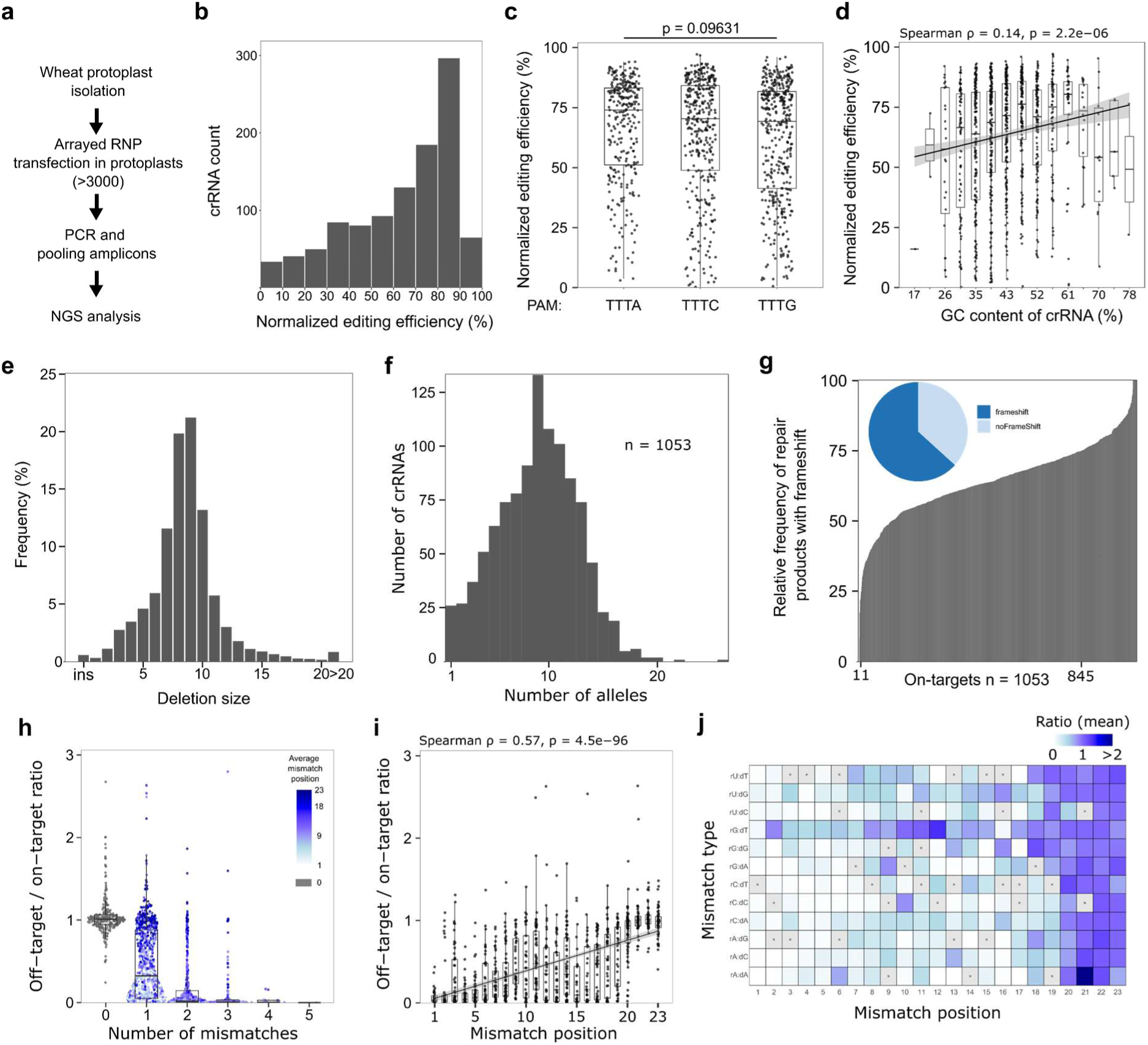
Arrayed CRISPR screen and Cas12a editing features. **(a)** Workflow of the arrayed CRISPR screen. 3000 RNPs were transfected in wheat protoplast and their activity was determined by NGS. **(b)** Histogram of normalized editing efficiency for LbCas12a crRNAs (n = 1061). **(c)** Impact of Cas12a PAM sequence on editing efficiency. Kruskal-Wallis p = 0.09631. **(d)** Impact of the crRNA GC content on editing efficiency. **(e)** Indel size distribution for Cas12a crRNAs. (**f)** Distribution of the number of alleles generated per crRNA (n=1053). **(g)** Distribution of indels with frameshift at individual target sites. Pie chart displays the proportion of indels resulting in frameshift mutations. **(h)** Off-target/on-target editing efficiency relative to the number of mismatches on the off-target site. Color scale indicates the average mismatch position for a given off target site. **(i)** Plotting off-target/on-target efficiency and the position of the mismatch for off-targets containing a single mismatch. **(j)** Heatmap displaying average off-target/on-target ratio for a given mismatch type and its position. Grey boxes with ‘x’ denote missing values.

### Editing efficiencies, repair outcomes and off-target analysis

The on-target editing efficiencies ranged from 0-100%, with skew towards values of >60% (**Fig. 2b**). We found no association between crRNA activity and the 4^th^ nucleotide in the PAM (**Fig. 2c**; p = 0.09631, One-way ANOVA) and only a very weak positive correlation with GC content (**Fig. 2d**; Spearman ρ = 0.14). The types of DNA repair products were consistent with LbCas12a activity; deletions were distributed around 8-9-bp and >1-bp insertions and >20-bp deletions occurred at low frequencies (**Fig. 2e**). Most crRNAs led to 10 or fewer unique alleles and approximately 1/3 of repair products were in-frame deletions (**Fig. 2f,g**). Between any two replicates, there is a strong correlation between specific allele frequencies (**Fig. S4a**), but there is also presence/absence variation; approximately one third of alleles are unique to a replicate and only 25% of alleles are found in all replicates (**Fig. S4b**). We estimate that each PCR had the potential to capture genotypes from as many as 3,000 cells. Taken altogether, while there are a limited number of possible alleles from any given crRNA, there also appears to be considerable variation between specific DNA repair products between replicates.

The number of mismatches between a crRNA and its putative target site(s) is an important feature when evaluating off-targets. Average off-target indel rates were negatively correlated with the number of mismatches (**Fig. 2h**). crRNAs with zero mismatches show very similar on- and off-target efficiencies, indicating that there does not appear to be a subgenomic effect. crRNAs with mismatches display reduced activity, with the positions being the controlling factor (**Fig. 2h,i**). Single mismatches along the spacer from positions 1-8 reduced average crRNA activity ∼75%, ∼50% from positions 9-18, and mismatches at positions 19-23 did not affect editing efficiency. Even for crRNAs with 2 or 3 mismatches, if these were located at the PAM-distal end, they still had appreciable activity. The type of mismatch has an effect as the three wobble mismatches (rG:dT, rU:dG, and rU:dT) led to average off-targeting rates of 55 – 78% as compared to other mismatches that only averaged 24 – 40% (**Fig. 2j, S5**).

The primary objective of the screen was to create a dataset to train a predictive model for LbCas12a crRNA activity in plants. Since several predictive models have already been reported in non-plant species, we decided to first compare our dataset to predicted scores from DeepCpf1. We observed a moderate Spearman correlation coefficient of 0.43 (**Fig. 3a**), which is considered to be at the lower bounds of being useful [20]. To bridge this gap, we decided to develop our own predictive model.

**Fig. 3:**
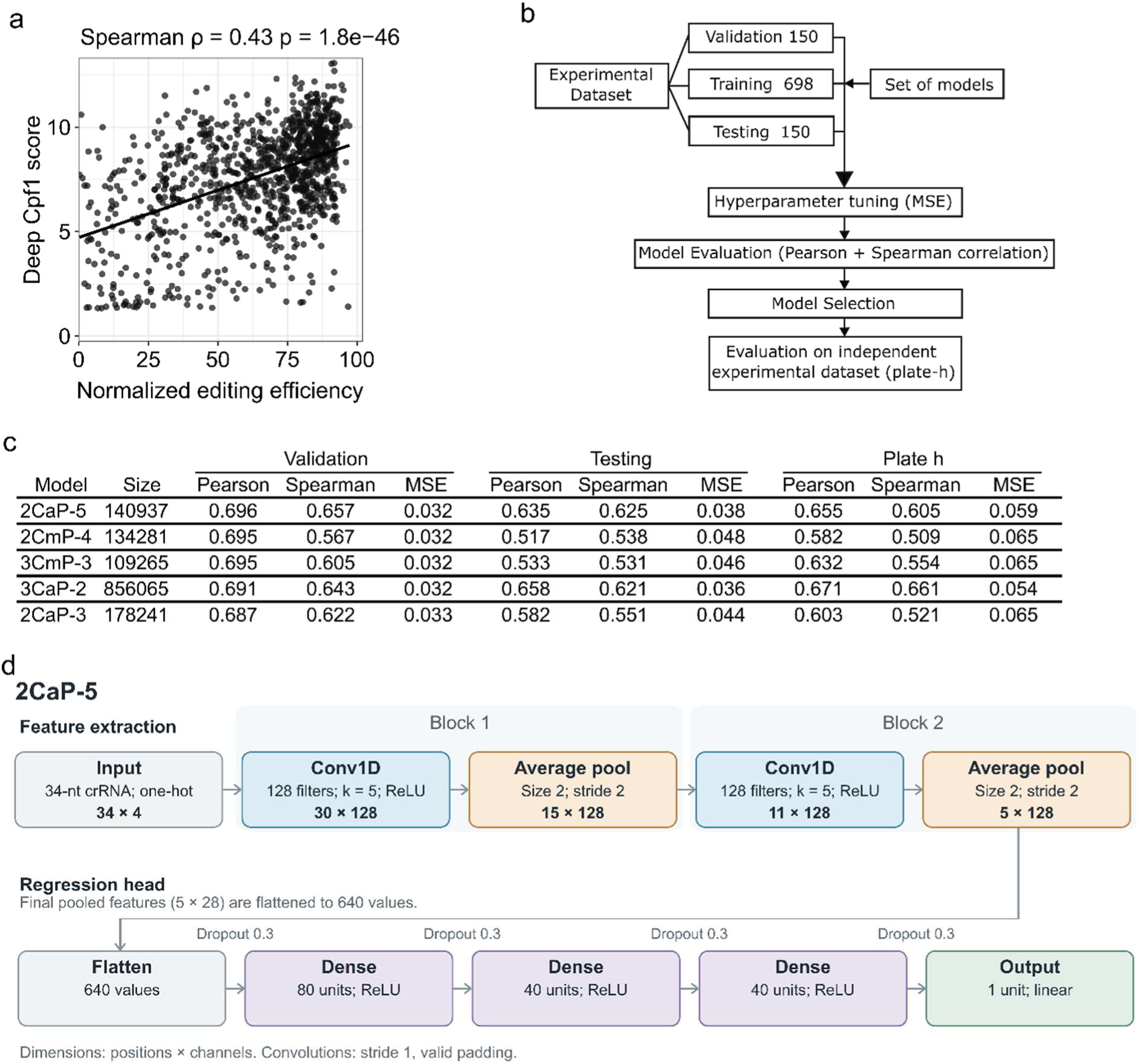
A machine learning model to predict LbCas12a activity. **(a)** Correlation between Deep-Cpf1 predictions from wheat crRNAs in this study and normalized editing efficiencies as measured by Spearman correlation (ρ=0.43). **(b)** Schematic representation of the machine learning model training and sub-division of datasets. Three independent datasets were used: validation and testing as part of the original screen and an independent dataset (plate-h) generated after the screen. **(c)** Model metrics for five models for validation, testing, and plate-h datasets. **(d)** Schematic representation of the 2CaP-5 model.

### Predictive modeling of Cas12a activity

From the total list of transfected crRNAs, we removed identical crRNAs and those that were eventually used for stable transformation (below) This left us with a set of 998 crRNAs and their efficiencies that were split into different datasets: training (698), validation (150) and testing (150). Five-fold cross-validation was done with the training set to identify the best hyperparameters (**Additional File 4 CNN and RNN results**). Subsequently, the model with the best hyperparameters was retrained on the entire training dataset and the validation data were used for early stopping. The testing data were kept separate to measure the generalization properties of the final chosen model. In addition to the testing dataset, we also performed an independent transfection experiment (plate h) with 91 unique crRNAs to provide a completely independent dataset to further benchmark the models.

Two ML architectures, Recurrent Neural Network (RNN) and Convolutional Neural Network (CNN), were used to generate a total of 40 distinct deep learning models to predict on-target editing efficiency and fine-tune the underlying hyperparameters (**Fig. 3b, Additional File 4 CNN and RNN results**). Mean Squared Error (MSE) was used as the cost function employed during training. The models were composed of two components: an extraction stack, responsible for capturing patterns from input sequences, and a regression stack, which employs the extracted patterns to compute the editing efficiency. The regression stack is composed of a dense neural network, while the extraction stack is tailored to various architectures (such as RNNs or 1D-convolutional layers combined with 1D-pooling layers). We adopted bi-directional RNNs to extract insights from both the 3’-5’ and 5’-3’ directions. However, these models required significantly longer training times, occasionally spanning several days to converge on the dataset. Contrastingly, CNN models exhibited both computational efficiency and rapid training convergence. The top models were all based on the CNN architecture (**Additional File 4 CNN and RNN results**).

A distinct model type, denoted as the ‘Base’ model, was also incorporated. This model adopts a standard Multilayer Perceptron configuration, where the extraction stack consists of a single dense layer (the bidirectional RNN layer or the 1D-convolution layer with a 1D-pooling layer (BiR/BiRd and CaP/CmP, respectively) were replaced with a conventional dense layer). The poor performance of the Base model compared to BiR/BiRd and CaP/CmP underscores the significance of positional information within DNA sequences for the prediction of editing efficiency (**Additional File 4 CNN and RNN results**). For similar reasons, CNNs with a kernel size of 1 also performed poorly.

Of the 40 architectures tested, 2CaP-5 had the best overall performance. It ranked first on the validation dataset and was within the top four on both the testing and plate-h datasets (**Fig. 3c**). This underscores the model’s robust generalization capabilities, reinforcing its effectiveness beyond the data used for training and validation sets. The 2CaP-5 model consists of two stacks of a 1D convolution layer with a kernel size of 5, each convolution layer followed by a 1D average pooling layer as a feature extractor (**Fig. 3d**). The feature extractor is followed by four linear layers each preceded by a dropout layer. The input consists of a one-hot encoding of the 34-bp DNA sequence comprising the 4 bp upstream of the PAM, the TTTV PAM, the 23 bp protospacer and 3 bp downstream. The model outputs a single scalar value between 0 and 1 which represents the predicted editing efficiency of the crRNA with Cas12a. The 2CaP-5 model is also relatively small, containing only ∼141,000 parameters (**Fig. 3d**, **Additional File 4 CNN and RNN results**), six-fold fewer than 3CaP-2, the only model with marginally higher correlation coefficients.

### Testing predicted crRNA efficiencies in wheat protoplasts and stable transformation

2CaP-5 was trained and validated on data from wheat protoplasts transfected with LbCas12a RNPs containing synthesized crRNAs. To determine if the model could be generalized to other experimental systems, we evaluated both transient expression in transfected wheat protoplasts and stable lines generated via Agrobacterium-mediated transformation. We compared 34 crRNAs that had a wide range of predicted (33-88%) and observed (6-92%) activity levels. The same LbCas12a and crRNA expression cassettes were used in both experimental systems: A maize Ubiquitin promoter controlled LbCas12a expression and a wheat U3 promoter was used to express crRNAs containing a truncated tRNA and two mature direct repeats [30]. For the protoplast transfections with plasmids, protoplasts were transfected and sorted via fluorescence activated cell sorting to enrich for cells expressing LbCas12a [30]. Mutation frequencies were determined by Illumina sequencing of the target sites and CRISPResso2 analysis. For the stably transformed plants, we obtained 3-14 plants for each of the six vectors, for a total of 54 plants. T0 leaf tissue was isolated for DNA extraction and genotyping was performed by Illumina sequencing and CRISPResso2 analysis. We obtained sequencing information from 92% of the on- and off-target loci. The allele frequencies were distributed in a clear W pattern, indicative of wild-type, heterozygous and biallelic genotypes (**Fig. S6a**). Per plant, the percentages of modified reads were distributed in a bimodal pattern with edited loci being predominantly biallelic and rarely heterozygous (**Fig. S6b**), indicating most mutagenesis occurred early during the transformation and/or tissue culture process.

When the 2CaP-5 predicted values of the 34 crRNAs are compared to the three experimental systems, we observe a moderate-to-strong correlation between the predicted values and both protoplast systems (**Fig. 4a-b**). However, there is no significant correlation between the 2CaP-5 predicted values and the efficiencies observed in plants (**Fig. 4c**). The bi-modal distribution of crRNA efficiencies in stable plants was not observed in any of the protoplast experiments where continuous values are observed. We do see a very strong correlation between the two protoplast transfection models (**Fig. 4d**). In contrast, we do not find a statistically significant correlation between the two protoplast-based systems and the stably transformed plants (**Fig. 4 e-f**).

**Fig. 4.**
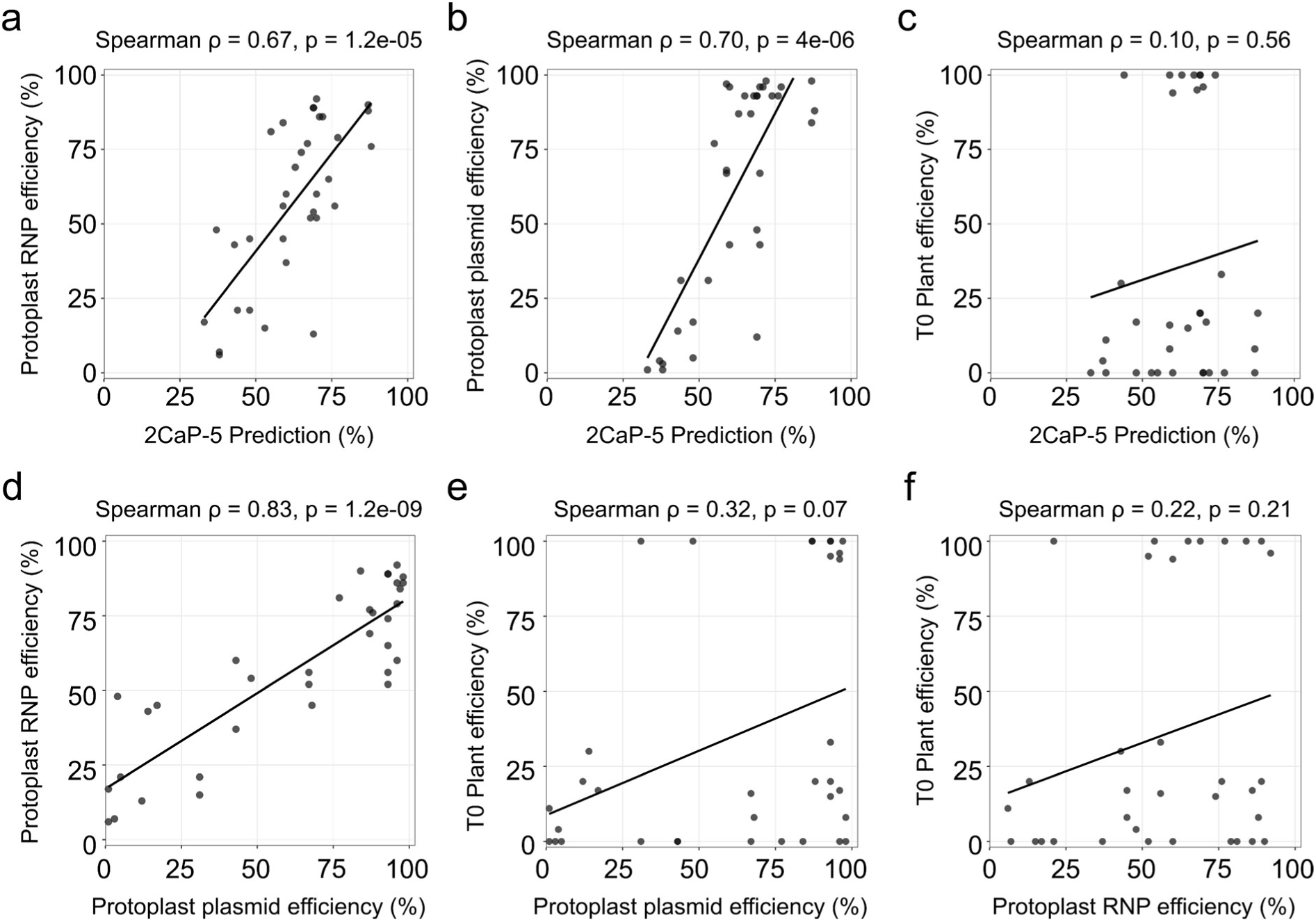
Correlations between different experimental systems. **(a)** 2CaP-5 predicted values vs protoplast RNP transfection, **(b)** 2CaP-5 predicted values vs. protoplast plasmid transfection, **(c)** 2CaP-5 predicted values vs. T0 plants, **(d)** protoplast plasmid transfection vs. crRNA, **(e)** protoplast plasmid transfection vs T0 plants, and **(f)** crRNA vs T0 plants. Spearman correlation coefficients ρ and exact p-values are indicated for each comparison. n = 34. T0 plant efficiency was calculated as the average proportion of mutated alleles.

### Correlation between DNA repair products

Our results indicated that protoplasts may not be an informative model for crRNA efficiencies in stably transformed plants. We wondered if there would still be some predictive power in regards to the types of mutant alleles generated. It is generally appreciated that DNA repair products are non-random and controlled by local DNA sequence contexts [3]. Reliably predicting which allele(s) might be produced would be valuable information prior to initiating transformation. We extracted all unique mutant alleles from the RNP, plasmid, and T0 plant experiments and calculated their relative abundances in each experiment. Similar to the comparison between RNP replicates (**Fig. S4**), only ∼half of the mutant alleles were shared between any two experimental systems for both on-targets and off-targets (**Fig. 5a-b**). If we only compare the shared alleles (minimum allele frequency ≥0.01), the relative abundances of unique alleles from RNP and plasmid experiments are strongly correlated (**Fig. 5c**). However, there is only a weak to moderate correlation between the protoplast models and the stable plants (**Fig. 5c-d**). For example, a 9-bp deletion was the most abundant allele for crRNA 296 in the RNP experiment and the second most abundant in the plasmid experiment. However, it was not detected in stably transformed plants. An 8-bp deletion was the most abundant allele from the plasmid experiment, but was only ranked 11^th^ in the RNP experiment and 2^nd^ in the stable plants (**Fig. 5e**). For crRNA 516, the three most abundant alleles are in agreement within the RNP and plasmid, one of which was the 2^nd^ most abundant in plants. Yet, the most abundant allele was ranked 6^th^ and 4^th^ for the RNP and plasmid experiments, respectively.

**Fig. 5.**
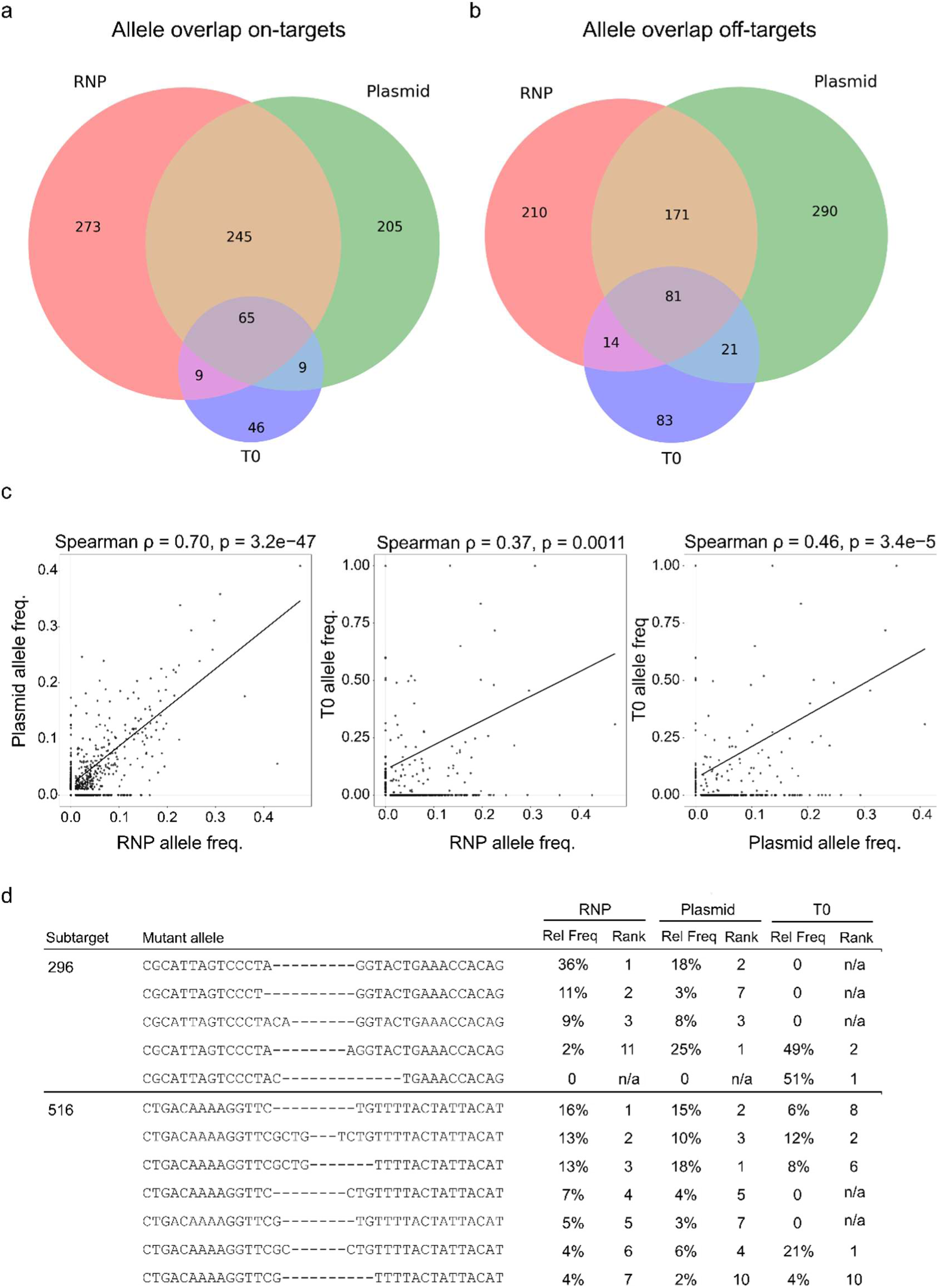
Reproducibility of DNA repair products across experimental systems. **(a)** Venn-diagram showing number of unique mutant alleles shared between RNP, Plasmid and T0 experiments for all on-targets and **(b)** off-targets. **(c)** Correlation of relative mutant allele frequencies between the different experimental systems. Alleles with at least 1% relative abundance are plotted. Spearman correlations are calculated on alleles that are shared between experimental systems. Samples with a zero value are omitted. **(d)** Representative examples of unique mutant alleles identified in the different experimental systems. For a given allele in a particular experimental system, the relative frequency (Rel Freq) and Rank are given. Only a subset of alleles are shown for clarity.

Altogether, these results indicate that the types of DNA repair products are moderately conserved across experimental systems. Correlations were stronger between protoplast-based assays than between protoplasts and stable transformation, further underscoring the limited predictive value of protoplast systems for repair outcomes in stably transformed plants.

### Predictive power of 2CaP-5 to small editing datasets

One possible explanation for the inability to predict crRNA efficiency in stable transformation is that the training data were based on protoplast data. If we want to develop a predictive model for stable plant experiments, these results suggest that stable plant data will most likely be necessary. Our training set contained efficiency scores from 698 unique crRNAs. While such numbers are on the low end to train deep-learning models, they are impractical for most stable plant transformation facilities. To determine the minimum number of guides one would need to create a reasonably effective predictive model, we performed a learning curve analysis on the 2CaP-5 model using different sizes of random subsets of the DeepCpf1 training dataset derived from human cell data [10]. Datasets containing just 100 crRNA sequences already improved the predictive power from a baseline Spearman correlation of ∼0.3 to ∼0.4 (**Fig.S7**). A maximum Spearman correlation of 0.74 was achieved when using 10,000 samples, which is similar to the fully trained DeepCpf1 model [10], but diminishing returns were already evident in datasets containing more than 500 crRNAs. Thus, we expect that data from at least 500- 1,000 crRNAs would be required to re-train 2CaP-5 from different experimental systems.

## Discussion

Plant genome editing experiments would greatly benefit from predictive models for target efficiency as many plant species require substantial time and resources to generate even tens of edited lines. As the number of gene targets become larger in scale with the use of multiplexing and CRISPR screens [22, 31], it becomes essential to generate high-efficiency reagents for robust and predictable experimental outcomes. This is especially important in asexual species as the regenerated genotype is fixed. Even in sexually reproducing species, increasing the number of target sites creates a significant downstream burden as the chances of finding homozygous or bi-allelic individuals at all target loci become exponentially smaller. Most work to date in plants has focused on improving the Cas enzyme or expression of the CRISPR reagents [32]. While several predictive models exist for Cas9 and Cas12a gRNAs and crRNAs, these models have been trained on mammalian data and have limited predictive power in plant systems (**Fig. 3a**; [14, 15]. As there is currently no available predictive model for gRNA or crRNA activity based on plant data, we aimed to bridge this gap by developing a crRNA predictive model trained on wheat protoplast data.

Starting with a dataset of only ∼1,000 crRNAs, we produced the 2CaP-5 model that is comparable with existing models [33]. The training data were derived from wheat protoplasts and we observed strong correlations between the model and activities in protoplasts. The format of the CRISPR reagents is apparently unimportant in protoplasts as RNP and plasmid transfections yielded highly similar results. However, these efficiencies did not translate to stable plants. Such discrepancies between experimental systems have been observed in mouse and human cell systems [13]. Therefore, researchers should exercise caution when extrapolating results from protoplast assays to stable transgenic plants, for instance when pre-testing crRNAs before stable transformation. While protoplast models are excellent to rapidly test the functionality of a given Cas12-crRNA editing machinery and establish genotyping assays, our data suggest that protoplasts may not provide quantitative insight into guide efficiency in plants stably transformed with *Agrobacterium*.

2CaP-5’s lack of predictive power for stably transformed plants is most likely caused by the low correlation between efficiencies in protoplast and stable plant systems. There are at least two main differences between protoplast transfection and stable plant transformation. One is the delivery of the reagents. For both RNP and plasmid transfections, millions of molecules are present per cell for a brief period of time (2 days). By contrast, *Agrobacterium* transformation probably delivers relatively few T-DNA molecules and, on average, only one or two are stably integrated, providing expression over a longer timeframe [34]. The second major difference is tissue type. Most protoplasts are derived from leaf mesophyll cells and do not divide whereas the immature embryos that are used as the explant for stable transformation undergo developmental reprogramming and cell division. If the transformation method (PEG vs. *Agrobacterium*) is the controlling factor, then a high-throughput *Agrobacterium* transient expression system could be utilized to generate a sufficient training set. If, however, the cell type is most important (protoplast vs. embryo), then large numbers of transformations would be required to obtain efficiencies for the 500-1,000 crRNAs needed to retrain 2CaP-5. With the use of multiplexing, the number of transformations could be reduced, but would still be on the order of hundreds. Additionally, sufficient plants should be obtained for every crRNA/transformation, ensuring that editing efficiencies can be reliably estimated. The fact that we were able to efficiently re-train 2CaP-5 on human cell line-derived data, indicates that 2CaP-5 can be generalized to other experimental systems.

Plant protoplasts are a widely used model system for various biological questions and technology development. Given that 2CaP-5 does work in wheat protoplasts, this model can already be used for a number of applications. While protocols to regenerate wheat plants from protoplasts are currently unavailable, a large number of plant species and genotypes can be regenerated from protoplasts [35, 36]. Following similar methods employed and the materials provided here, researchers could obtain data from 500-1000 crRNAs/gRNAs to re-train 2CaP-5 for their specific species of interest.

From our panel of crRNAs in wheat we can derive some general LbCas12a indel features at on- and off-target sites. Generally, crRNAs with a GC content ∼30-70% are efficient at inducing deletions and the majority of LbCas12a deletions are between 7 and 11-bp in length. Such larger deletions have been reported in various LbCas12a experiments in plants and other systems [3, 37, 38]. Interestingly, the chance of creating an in-frame mutation is approximately one-in-three. This stands in contrast to Cas9-based indels, which are typically ±1bp and less frequently result in in-frame mutations [39]. This feature of LbCas12a could be particularly useful when creating protein or promoter variants rather than gene KOs [40].

Our dataset also provides an extensive analysis of LbCas12a off-target activity in a plant system. We find that LbCas12a is highly sensitive to single nucleotide mismatches within the first 8 nucleotides from the PAM and even mismatches up to position 18 can decrease average activity by more than 50%. Overall, off-targets with ≥3 mismatches result in little to no detectable activity. Off-targets that do maintain activity tend to have mismatches in the PAM-distal region. Interestingly, mismatch identity is an important design criterium as RNA:DNA mismatch wobbles effectively doubled the rate of off-targeting compared to other mismatch types. These findings provide practical on- and off-target design guidelines for crRNA selection in highly homologous genomic contexts. Specifically, introducing two to three non-wobble mismatches within the PAM-proximal region (positions 1–8) should effectively minimize off-target activity. Conversely, when targeting homeologous genes, mismatches should be preferentially positioned in the PAM-distal region of the crRNA to retain editing efficiency and ideally be composed of RNA:DNA wobble pairs. Note that rG:dT, rU:dG, and rU:dT mismatches correspond to the spacer:target pairs whereas crRNA design compares spacer sequences. When designing crRNAs, these would correspond to G:A, T:C, and T:A mismatches, respectively.

An unexpected observation is that the LbCas12a editing outcomes varied widely between experimental systems as only ∼half of the observed alleles were shared between experimental systems and independent replicates of RNPs. This stands in contrast to Cas9 which tends to produce very specific, and reproducible, 1-bp insertions [3, 41]. A possible explanation could be limited sample sizes. If our DNA extraction efficiencies are 100%, then we would expect the equivalent of 3,000 genomes, or 6,000 unique alleles per subtarget in the PCRs used for genotyping our RNP experiments. DNA extractions are not 100%, but we could still expect capturing hundreds of alleles per replicate. Furthermore, we did not observe such variation in the indel efficiencies. In stark contrast to the indel type, indel frequencies were highly reproducible between replicates in protoplast systems. This suggests that the variations in specific DNA repair outcomes are indeed real observations and warrant follow-up investigations.

## Conclusions

Here, we report a large-scale CRISPR screen in wheat protoplasts to develop a predictive model for crRNA efficiency. Our 2CaP-5 model provides a strong foundation for researchers using LbCas12a in wheat and potentially other plant protoplasts systems. We demonstrate that 2CaP-5 can be efficiently retrained, making it a generally useful tool for the (plant) genome editing community. Our extensive off-target analysis also reinforces the specificity of LbCas12a systems, especially with regards to the number and type of mismatches and their relative position within the crRNA.

## Material & Methods

### Plant growth conditions

The wheat cultivar “Fielder” was used in all experiments. Two batches of seeds (7.5 g) were sterilized by successive washes with sterilized water for 3 min, isopropanol for 45 sec, sterile water for 3 min and 6% sodium hypochlorite (Chem-lab nv) for 10 min, and six water washes before sowing. Seeds were grown on 1/2 MS pH 5.7 (Duchefa Biochemie, M0221.0050), 2.5 mM MES (M1503.0100, Duchefa Biochemie) and 0.5% plant tissue agar (NCM0250A, NEOGEN) in cylindrical containers (960162, Greiner Bio-one). Per batch, 160 seeds were sown in 80 cylinders and stratified at 4°C in the dark for 3 or 4 days. Plants were grown under SpectraluxPlus NL 36 W/ 840 Plus (Radium Lampenwerk) fluorescent bulbs under long days (16h light/8h dark) for 8 days at 25°C.

### Ribonucleic Protein Complexes (RNP) assembly

Individual RNPs were assembled by mixing 0.6 µl LbCas12a ultra v1 or v2 at 10 µg/µl (Alt-R™ L.b.Cas12a (Cpf1) Ultra (IDT)), 0.8 µl crRNAs (96 well Alt-R CRISPR Plate, IDT) in Duplex buffer at 100 µM, 0.55 µl NEB 2.1 buffer and 3.55 µl RNAse-free water. A master mix of LbCas12a, NEB buffer and water was first distributed into a 96-well plate and crRNAs were individually added using a multi-channel pipette. Reagents were then briefly mixed and incubated at room temperature for 25 min. Samples were kept on ice until protoplast transfections. 12 µg of pZmUBI-GFP-NLS-G7T was used as a carrier plasmid.

### Protoplast isolation and transfection

Protoplast isolation and RNP or plasmid transfection were conducted similarly as described before [30], with minor changes. Wheat leaves were harvested 7 days after germination. Two batches of 45 leaves were cut in 0.5-1 mm strips before incubation in 0.6 M D-mannitol (M1902, Sigma-Aldrich) for 10 min in the dark. Samples were successively incubated in enzyme solution (20 mM MES, 1.5% cellulase R10 (C8001.0010, Duchefa Biochemie), 0.75% macerozyme R10 (M8002.0010, Duchefa Biochemie), 0.6 M D-mannitol and 10 mM KCl, 0.1% BSA and 10 mM CaCl_2_) for 8 hours, filtered through a 40 µm cell strainer (#431750, Corning), collected by centrifugation at 80×g and incubated in W5 solution (2 mM MES pH 5.7, 154 mM NaCl, 125 mM CaCl_2_ 0.5 mM KCl) for 30 min. Protoplasts were then counted with a Neubauer chamber and adjusted to a concentration of 1 × 10^6^ cells ml^-1^ in MMG solution (4 mM MES pH 5.7, 0.4 M mannitol, 15 mM MgCl_2_). The two independent batches were combined and incubated on ice for 30 min before transfection.

Three RNPs were mixed (3 × 5.5 µl) and 3.5 µl MMG solution was added to 1 ml strip tubes (TN0946-08B, National Scientific Supply Co) and kept on ice. In total, 96 x 3 RNPs were distributed according to the pattern of three individual crRNA plates (**Fig. S3a**). 100 μl of protoplasts (10^5^ cells) and 110 μl of PEG solution (0.2 M mannitol, 100 mM CaCl_2_), 40% PEG 4000 (81240, Sigma) were added to RNPs or plasmid DNA using a multichannel pipette and immediately mixed by slowly inverting the strip. Individual strips (from 1 to 12) were transfected at a fixed time interval (1.5 min) and incubated for 18 min. W5 was added at the same time interval and strips were inverted 4-5 times to stop the transfection reaction. Protoplasts were then collected after centrifugation at 80×g, diluted in 900 µl W5 and incubated at 25 °C in dark for 44 hours. Wheat protoplasts were harvested by centrifugation at 4000×g for 3 min and removal of the supernatant. Samples were frozen in liquid nitrogen and stored at -20 °C.

DNA was isolated according to the Edwards method [42]. Two metal beads (3 mm) were added, and tissue was ground by shaking the plate by hand. 400 µl of extraction buffer (100 mM Tris-HCl pH 8.0, 500 mM NaCl, 50 mM EDTA, 0.7% SDS) was added to individual samples and incubated for 30 min at 60 °C. Samples were centrifuged for 5 minutes and 300 µl of the supernatant was transferred to a new tube. The supernatant was mixed with 300 µl isopropanol for DNA precipitation. After centrifugation, the pellet was washed with 70% Ethanol, dried at room temperature, and dissolved in 100 µl of 10 mM Tris-HCl pH 8.0. 50 µl of the samples were transferred to a 96-well PCR plate and the remaining solution was kept as stock in -20 °C.

### Genotyping

The amplicons used to establish the screening conditions were designed using primer pairs amplifying 600 bp around the TaMLO-D, TS13-3, TS60-3, TS92-1, TS196-3, TS230-1 target sites. For the MLO internal control, unique pairs of index primers amplifying a ∼250 bp region were used (**Additional File 2 Primers**). For all other target sites, unique primer pairs were designed to amplify a region ranging from 170-250 bp. All primers were ordered from Integrated DNA Technologies (IDT) and amplicons for all target sites were obtained by PCR using Q5® High-Fidelity DNA Polymerase (M0491, NEB) using 1 µl of gDNA as template. PCR products were purified using magnetic beads (AC-60500, MAGBIO Genomics), eluted in 35 μl 10 mM Tris-HCl pH 8.0 and sent for Sanger sequencing (Mix2Seq, Eurofins). Trace files of the chromatograms were analyzed using ICE (Synthego [29]).

The CRISPR screen amplicons were generated using the following PCR conditions: initial 98 °C: 30s, 30 cycles of 98 °C for 10s, 62 °C for 20s, 72 °C for 20s; final 72 °C for 5 min. 4 µl of individual PCR samples were pooled and column purified using the PCR Purification Kit (D4013, Zymo Research Co.). Pooled DNA was run on 2% agarose gel (Ultra-Pure agarose, # 16500-500, Invitrogen), with a Low molecular Weight DNA ladder (NEB, B7025) and PCR products around 200-250 bp were gel purified (D4002, Zymo Research). Pooled DNA concentration was then measured with Qubit (Invitrogen) according to manufacturer’s protocol and adjusted to 2 ng µl^-1^. Paired end sequencing was performed with Eurofins NGSelect amplicons (5M reads with Illumina PE150).

All primers, amplicons, target sites and crRNA used in this study are found in the Supplemental information.

### Illumina sequencing and statistical analysis

Editing efficiencies from Illumina sequencing files were obtained with CRISPResso2 [43] via a Galaxy workflow (https://usegalaxy.be/) (**Additional File 5 Demultiplex CRIPSR screen and Additional File 6 CRISPResso Batch CRISPR screen**). Sequence files (fastq.gz) together with a description file containing the name and sequence of all three wheat subgenome amplicons and crRNAs were used. Cleavage offset was set to -4, with default editing window parameter and minimum average read quality to 30. Separate files for individual target sites were concatenated. During analysis we found the -4 setting missed a number of legitimate edited alleles. We therefore converted all ‘Unedited’ cells in the CRISPResso2 output to FALSE if the ‘Aligned_Sequence’ contained a dash ‘-‘.

For MLO amplicons, reads were demultiplexed using Je-demultiplex with a barcode length of 15 and individual fastq files were obtained with a Galaxy workflow as previously described [30]. Individual fastq files were generated for MLOA and MLOBD. CRISPRessoBatch was used to analyze MLOA and CRIPSRessoPooled was used for MLOB and MLOD. Individual editing result files were then combined and the unedited column was corrected as described above.

For the establishment of the screening method using TS13-3 and MLO-D, the “modified” data were treated as binary outcomes using a Generalized Linear Model with a logit link function *g*(*µ*), assuming a Binomial distribution. Two GLMs were fitted to estimate replicate, row and column effects.

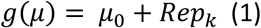

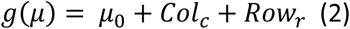

Model 1 was fitted to all plates and allows the estimation of replicate effects, while model 2 was fitted to the plates separately and estimates the row and column effects on a single plate. The overall significances of the fixed plate, row and column effects were assessed by a likelihood test for dropping fixed terms, as implemented in Genstat (Genstat for Windows 22^nd^ Edition. VSN International, Hemel Hempstead, UK. Web page: Genstat.co.uk). T statistics were used to assess the significance of specific replicate, column or row effects by pairwise comparisons (on the logit scale) to the reference level.

For the RNP screen, the data were treated the same except a Generalized Linear Mixed Model, was fitted, with a logit link function *g*(*µ*), assuming a Binomial distribution for the fixed effects and a Beta distribution for the random effects. Two GLMMs were fitted to model batch effects and transfection and amplicon as random effects.

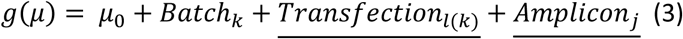

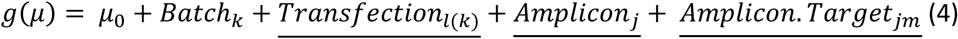

Model 3 was fitted to all three batches, while model 4 was fitted to only two batches corresponding with Targets GenomeA and GenomeB. For both models, the dispersion parameter for the variance of the response was estimated from the residual mean square of the fitted model. Combined indel efficiencies were then normalized using the Logit normalization method based on the efficiencies of MLO RNPs distributed on all plates. All other statistical analyses were done using GraphPad Prism v9.0.0.121 or higher and R v3.5.1.

T0 wheat editing efficiencies were obtained by averaging the proportion of mutated alleles per target. Raw values can be found in **Additional File 7 T0 genotyping**.

### Target sites, primers and crRNAs designs

We identified 1,639 wheat homeologous triplets (4,917 genes) and selected one homeolog of each triplet at random to be the target, while the two homeologs became the off-targets. We retrieved the full length genomic sequence from the Chinese Spring reference genome (including introns and UTRs) of the selected target genes and cut the target sequences into 250 bp parts (size of amplicon). We obtained 26,669 possible amplicons and used BLAST against a proprietary Fielder reference genome (BASF) to retain only those with a 100% full length match to Fielder. We removed amplicons with a GC content of less than 30% or over 70%. To avoid guides at the edge of the sequences where primers would bind, we removed 75 bp at both ends, leaving a 100 bp window in the center for guide design. 20,995 potential amplicons remained and within this set we found 53,454 possible TTTVN23 Cas12a target sites.

We filtered the results to only get crRNAs that have exactly three targets, corresponding to one target gene with 0 mismatches and two off-target genes with at least 1 mismatch, where the target and off-target genes constituting a homeolog triplet. We removed any guides with GC% under 30% or over 80%. We selected 1,000 crRNAs with the following parameters: 40% in CDS, 60% different mismatches between off-targets, 40% with 1 mismatch to both off-target and 40% with at most 2 mismatches.

### Primer design

We aligned all three amplicons with MUSCLE [44] and created a stringent consensus sequence to design primer pairs to amplify all three subgenomes. We used primer3 [45] to design primers on the consensus sequences. We specified the positions of the left and right primers so that they are not too close to the crRNA and not too close to the ends of the amplicon and ask for the best primer pair to be returned. We selected 1,056 unique crRNAs with corresponding primer pairs for 1056 × 3 amplicons. Primers were ordered on 12 independent plates (IDT) and were used to assess correct PCR amplification and detection with CRISPResso2 [43].

### Cloning

Vectors for stable wheat transformation were constructed by combining several cassettes (crRNA array, LbCas12a, linkers, TaGRF4-TaGIF1 morphogenic regulator, hygromycin selection) using Golden Gate assembly followed by Gibson assembly in the pG3-U1-AG-U9 vector backbone [46, 47] compatible with the ternary vector pVS1-VIR2 (Addgene #134745, [48]).

Entry clones for six crRNA cassettes were built by Golden Gate assembly of annealed oligos in vectors containing a crRNA expression cassette (Additional file Plasmids). Each array contained 6 crRNAs driven by a wheat U3 promoter and was assembled by Golden Gate assembly with BsaI using the following program (30 cycles (37°C, 5 min; 16°C, 5 min); 50°C for 5 min; 80°C for 5 min). The two crRNAs 359 and 509 were removed from later analyses as we could not distinguish on-and off-target reads in mutated samples (ambiguous targets). The LbCas12a cassette was built by Golden Gate assembly with the ZmUbi promoter and G7 terminator (**Additional File 8 Plasmids**). The pGGIB-U4-pPvUBI2-TaGRF4-TaGIF1-tBdUBI1-U5, pGGIB-U8-pOsAct-HI-t35S-U9 and linker cassettes were previously described [46]. All entry cassettes were digested for 2 hours by I-SceI and assembled by a Gibson reaction at 50°C for 2 hours. Entry vectors were quality-checked using Sanger sequencing (Mix2Seq, Eurofins) and binary vectors were verified by restriction digestion followed by whole plasmid-sequencing using Oxford Nanopore long read sequencing (PlasmidSaurus). All plasmids are deposited on (https://vectorvault.vib.be/).

### Machine learning model

The dataset designed for on-target predictions, consisting of 998 distinct crRNA sequences and their corresponding efficiencies, underwent a division into 698 training samples, 150 validation samples, and 150 testing samples. During the model’s training, hyperparameter optimization, and final model selection stages, the training and validation sets were solely employed. The testing dataset, in conjunction with an independent experiments dataset (**Additional File 9 Plate-h dataset**), was harnessed to gauge the ultimate model’s generalizability. No overlap in terms of crRNA sequences existed between any of the datasets.

Before initiating the model training, crRNA sequences underwent tokenization and were subsequently one-hot encoded. Training proceeded with the inclusion of early stopping mechanisms, with a patience of 25 epochs. The hyperparameters and their respective values associated with each model can be found within **Additional File 4 CNN and RNN results**.

All model training was executed on the UGent High-Performance Computing (HPC) infrastructure. Some models were trained on the Joltik cluster, equipped with 2x 16-core Intel Xeon Gold 6242 processors (Cascade Lake @ 2.8 GHz) and 4x NVIDIA Volta V100 GPUs (32GB GPU memory). Others were trained on the Accelgor cluster, featuring 2x 24-core AMD EPYC 7413 processors (Milan @ 2.2 GHz) and 4x NVIDIA Ampere A100 GPUs (80GB GPU memory). TensorFlow version 2.11.0 served as the framework for model development and training.

### Stable wheat transformation

*Agrobacterium*-mediated wheat transformation was performed as previously described [49], with some modifications. The *Agrobacterium* strain EHA105 *recA*-containing pVS1-VIR2 was transformed with the binary vectors using the freeze-thaw method [50]. Clones were verified using colony PCR. *Agrobacterium* cells were streaked on YP medium (12 g L^-1^ yeast extract, 10 g L^-1^ peptone, 5 g L^-1^ NaCl, pH 6.8, 8 g L^-1^ agar) supplemented with antibiotics and grown for 3 days at 24 °C. One day before transformation, a loop of the bacteria was resuspended in 200 µl sterile H_2_O, plated on a fresh YP plate and incubated at 28 °C overnight. A loop of overnight *Agrobacterium* culture was mixed with 5 ml wheat inoculation medium + 100 µM acetosyringone (AS) and shaken for 3-4 hours at 200 rpm.

Wheat donor plants were grown in a growth cabinet at 19 °C with a 16h photoperiod. Six to ten flowering ears were used per construct and experiment. Calli and regenerating shoots were grown under SpectraluxPlus NL 36 W/ 840 Plus (Radium Lampenwerk) fluorescent bulbs under long days (16h light/8h dark) at 25°C. Green shoots arising from the calli were selected as transgenic events. Multiple shoots arising from individual calli were considered clonal. Green shoots from single plantlets were transferred to rooting medium for 2-3 weeks. Plant leaf tissue was collected for genotyping and frozen in liquid nitrogen and the plantlets were transferred to a soilless potting mix for propagation.

## Supporting information

Additional File 1: All amplicons

Additional File 2: Primers

Additional File 3: crRNAs

Additional File 4: CNN and RNN results

Additional File 5 CRISPResso Batch CRISPR screen

Additional File 6 Demultiplex CRIPSR screen

Additional File 7: T0 genotyping

Additional File 8: Plasmids

Additional File 9: Plate-h dataset

Additional File 10: Raw data file

## Additional Files

Additional File 1: All amplicons

Additional File 2: Primers

Additional File 3: crRNAs

Additional File 4: CNN and RNN results

Additional File 5: CRISPResso Batch Galaxy pipeline settings

Additional File 6 Demultiplex CRIPSR screen Galaxy pipeline settings

Additional File 7: T0 genotyping

Additional File 8: Plasmids

Additional File 9: Plate-h dataset

Additional File 10: Raw data file

Additional File 11: Supplemental figures. **Fig. S1**: Validation primer pair for target site amplicons. **Fig. S2**. Establishment of 96-well RNP transfection and indel frequency variability with transfection replicate and plate position. **Fig. S3.** CRISPR screen statistics. **Fig. S4**. Allele overlap between RNP replicates. **Fig. S5**. RNA:DNA mismatch type affects off-targeting rates. **Fig. S6**. T0 plant allele frequencies distribution. **Fig. S7**. 2CaP-5 learning curve.

## Declarations

### Ethics approval and consent to participate

Not applicable

### Consent for publication

Not applicable

### Availability of data and materials

All data supporting the findings of this study are available within the paper and its Supplementary Information. All raw data can be found in Additional File 10 Raw data. The 2CaP-5 model and associated analyses are available in the GitHub repository https://github.com/VIB-PSB/crRNA-Predictive-Modeling.

### Authors’ contributions

CG: crRNA and screen design, RNP optimization, Protoplasts transfection and processing, NGS analysis, stable plant transformation, manuscript preparation. PW: Computational modeling, manuscript preparation. DM: computational modeling. LS: NGS analysis and figure preparation. MS: crRNA design, NGS analysis. DDV: Screen design, RNP optimization, protoplast transfection. MV: statistical analysis. APF: Protoplasts transfection and processing, stable wheat transformation. JdS: Protoplast transfection. DR, EVL, TZ, IDS, LP: Stable wheat transformation and genotyping. WD: NGS analysis. ES: NGS analysis, manuscript preparation. DI: Computational modeling, manuscript preparation. FM, TBJ: Project lead, crRNA and screen design, NGS analysis, manuscript preparation. All authors read and approved the final manuscript.

## Acknowledgements

We thank Sadiye Hayta, Mark A. Smedley and Wendy A. Harwood for training in wheat transformation.

## References

1. Paul B, Montoya G: CRISPR-Cas12a: Functional overview and applications. BIOMEDICAL JOURNAL 2020, 43:8–17.

2. Kock MA: Open Intellectual Property Models for Plant Innovations in the Context of New Breeding Technologies. In Agronomy, vol. 11. pp. 1218; 2021:1218.

3. Allen F, Crepaldi L, Alsinet C, Strong AJ, Kleshchevnikov V, De Angeli P, Palenikova P, Khodak A, Kiselev V, Kosicki M, et al: Predicting the mutations generated by repair of Cas9-induced double-strand breaks. Nature Biotechnology 2019, 37:64-+.

4. Van Huffel K, Stock M, Ruttink T, De Baets B: Covering the Combinatorial Design Space of Multiplex CRISPR/Cas Experiments in Plants. Frontiers in Plant Science 2022, 13.

5. Doench JG, Fusi N, Sullender M, Hegde M, Vaimberg EW, Donovan KF, Smith I, Tothova Z, Wilen C, Orchard R, et al: Optimized sgRNA design to maximize activity and minimize off-target effects of CRISPR-Cas9. Nature Biotechnology 2016, 34:184-+.

6. Shen M, Arbab M, Hsu J, Worstell D, Culbertson S, Krabbe O, Cassa C, Liu D, Gifford D, Sherwood R: Predictable and precise template-free CRISPR editing of pathogenic variants. NATURE 2018, 563:646-+.

7. Kim H, Kim Y, Lee S, Min S, Bae J, Choi J, Park J, Jung D, Yoon S, Kim H: SpCas9 activity prediction by DeepSpCas9, a deep learning-based model with high generalization performance. SCIENCE ADVANCES 2019, 5.

8. Kim N, Kim H, Lee S, Seo J, Choi J, Park J, Min S, Yoon S, Cho S, Kim H: Prediction of the sequence-specific cleavage activity of Cas9 variants. NATURE BIOTECHNOLOGY 2020, 38:1328-+.

9. Kim H, Song M, Lee J, Menon A, Jung S, Kang Y, Choi J, Woo E, Koh H, Nam J, Kim H: In vivo high-throughput profiling of CRISPR-Cpf1 activity. NATURE METHODS 2017, 14:153–159.

10. Kim H, Min S, Song M, Jung S, Choi J, Kim Y, Lee S, Yoon S, Kim H: Deep learning improves prediction of CRISPR-Cpf1 guide RNA activity. NATURE BIOTECHNOLOGY 2018, 36:239-+.

11. Zhu H, Liang C: CRISPR-DT: designing gRNAs for the CRISPR-Cpf1 system with improved target efficiency and specificity. BIOINFORMATICS 2019, 35:2783–2789.

12. O’Brien A, Bauer D, Burgio G: Predicting CRISPR-Cas12a guide efficiency for targeting using machine learning. PLOS ONE 2023, 18.

13. Haeussler M, Schonig K, Eckert H, Eschstruth A, Mianne J, Renaud JB, Schneider-Maunoury S, Shkumatava A, Teboul L, Kent J, et al: Evaluation of off-target and on-target scoring algorithms and integration into the guide RNA selection tool CRISPOR. Genome Biology 2016, 17:12.

14. Naim F, Shand K, Hayashi S, O’Brien M, McGree J, Johnson A, Dugdale B, Waterhouse P: Are the current gRNA ranking prediction algorithms useful for genome editing in plants? PLOS ONE 2020, 15.

15. Slaman E, Lammers M, Angenent G, de Maagd R: High-throughput sgRNA testing reveals rules for Cas9 specificity and DNA repair in tomato cells. FRONTIERS IN GENOME EDITING 2023, 5.

16. Slaman E, Kottenhagen L, de Martines W, Angenent G, de Maagd R: Comparison of Cas12a and Cas9-mediated mutagenesis in tomato cells. SCIENTIFIC REPORTS 2024, 14.

17. Li S, Zhang C, Li J, Yan L, Wang N, Xia L: Present and future prospects for wheat improvement through genome editing and advanced technologies. PLANT COMMUNICATIONS 2021, 2.

18. Athiyannan N, Abrouk M, Boshoff W, Cauet S, Rodde N, Kudrna D, Mohammed N, Bettgenhaeuser J, Botha K, Derman S, et al: Long-read genome sequencing of bread wheat facilitates disease resistance gene cloning. NATURE GENETICS 2022, 54:227-+.

19. Walkowiak S, Gao L, Monat C, Haberer G, Kassa M, Brinton J, Ramirez-Gonzalez R, Kolodziej M, Delorean E, Thambugala D, et al: Multiple wheat genomes reveal global variation in modern breeding. NATURE 2020, 588.

20. Konstantakos V, Nentidis A, Krithara A, Paliouras G: CRISPR-Cas9 gRNA efficiency prediction: an overview of predictive tools and the role of deep learning. NUCLEIC ACIDS RESEARCH 2022, 50:3616–3637.

21. LeCun Y, Bengio Y, Hinton G: Deep learning. NATURE 2015, 521:436–444.

22. Hu YJ, Patra P, Pisanty O, Shafir A, Belew ZM, Binenbaum J, Ben Yaakov S, Shi BH, Charrier L, Hyams G, et al: Multi-Knock-a multi-targeted genome-scale CRISPR toolbox to overcome functional redundancy in plants. Nature Plants 2023, 9:572-+.

23. Liu H, Jian L, Xu J, Zhang Q, Zhang M, Jin M, Peng Y, Yan J, Han B, Liu J, et al: High-Throughput CRISPR/Cas9 Mutagenesis Streamlines Trait Gene Identification in Maize. PLANT CELL 2020, 32:1397–1413.

24. Lu Y, Ye X, Guo R, Huang J, Wang W, Tang J, Tan L, Zhu J-k, Chu C, Qian Y: Genome-wide Targeted Mutagenesis in Rice Using the CRISPR/Cas9 System. Molecular Plant 2017.

25. Meng X, Yu H, Zhang Y, Zhuang F, Song X, Gao S, Gao C, Li J: Construction of a Genome-Wide Mutant Library in Rice Using CRISPR/Cas9. MOLECULAR PLANT 2017, 10:1238–1241.

26. Berman A, Su N, Li Z, Landau U, Chakraborty J, Gerbi N, Liu J, Qin Y, Yuan B, Wei W, et al: Construction of multi-targeted CRISPR libraries in tomato to overcome functional redundancy at genome-scale level. NATURE COMMUNICATIONS 2025, 16.

27. Sun L, Alariqi M, Wang Y, Wang Q, Xu Z, Zafar M, Yang G, Jia R, Hussain A, Chen Y, et al: Construction of Host Plant Insect-Resistance Mutant Library by High-Throughput CRISPR/Cas9 System and Identification of A Broad-Spectrum Insect Resistance Gene. ADVANCED SCIENCE 2024, 11.

28. Woo JW, Kim J, Il Kwon S, Corvalan C, Cho SW, Kim H, Kim SG, Kim ST, Choe S, Kim JS: DNA-free genome editing in plants with preassembled CRISPR-Cas9 ribonucleoproteins. Nature Biotechnology 2015, 33:1162–U1156.

29. Conant D, Hsiau T, Rossi N, Oki J, Maures T, Waite K, Yang JY, Joshi S, Kelso R, Holden K, et al: Inference of CRISPR Edits from Sanger Trace Data. Crispr Journal 2022, 5:123–130.

30. Gaillochet C, Fernandez AP, Goossens V, D’Halluin K, Drozdzecki A, Shafie M, Van Duyse J, Van Isterdael G, Gonzalez C, Vermeersch M, et al: Systematic optimization of Cas12a base editors in wheat and maize using the ITER platform. Genome Biology 2023, 24.

31. Lorenzo CD, Debray K, Herwegh D, Develtere W, Impens L, Schaumont D, Vandeputte W, Aesaert S, Coussens G, De Boe Y, et al: BREEDIT: a multiplex genome editing strategy to improve complex quantitative traits in maize (vol 35, 218, 2023). Plant Cell 2023.

32. Zhou J, Luan X, Liu Y, Wang L, Wang J, Yang S, Liu S, Zhang J, Liu H, Yao D: Strategies and Methods for Improving the Efficiency of CRISPR/Cas9 Gene Editing in Plant Molecular Breeding. PLANTS-BASEL 2023, 12.

33. Yuan H, Song C, Xu H, Sun Y, Anthon C, Bolund L, Lin L, Benabdellah K, Lee C, Hou Y, et al: An Overview and Comparative Analysis of CRISPR-SpCas9 gRNA Activity Prediction Tools. CRISPR JOURNAL 2025, 8:89–104.

34. Oltmanns H, Frame B, Lee L, Johnson S, Li B, Wang K, Gelvin S: Generation of Backbone-Free, Low Transgene Copy Plants by Launching T-DNA from the Agrobacterium Chromosome. PLANT PHYSIOLOGY 2010, 152:1158–1166.

35. Reyna-Llorens I, Ferro-Costa M, Burgess S: Plant protoplasts in the age of synthetic biology. JOURNAL OF EXPERIMENTAL BOTANY 2023, 74:3821–3832.

36. Reed K, Bargmann B: Protoplast Regeneration and Its Use in New Plant Breeding Technologies. FRONTIERS IN GENOME EDITING 2021, 3.

37. Leenay R, Aghazadeh A, Hiatt J, Tse D, Roth T, Apathy R, Shifrut E, Hultquist J, Krogan N, Wu Z, et al: Large dataset enables prediction of repair after CRISPR-Cas9 editing in primary T cells. NATURE BIOTECHNOLOGY 2019, 37:1034-+.

38. Kleinstiver B, Tsai S, Prew M, Nguyen N, Welch M, Lopez J, McCaw Z, Aryee M, Joung J: Genome-wide specificities of CRISPR-Cas Cpf1 nucleases in human cells. NATURE BIOTECHNOLOGY 2016, 34:869-+.

39. Chakrabarti A, Henser-Brownhill T, Monserrat J, Poetsch A, Luscombe N, Scaffidi P: Target-Specific Precision of CRISPR-Mediated Genome Editing. MOLECULAR CELL 2019, 73:699-+.

40. Zhou J, Liu G, Zhao Y, Zhang R, Tang X, Li L, Jia X, Guo Y, Wu Y, Han Y, et al: An efficient CRISPR-Cas12a promoter editing system for crop improvement. NATURE PLANTS 2023, 9:588-+.

41. Weiss T, Kumar J, Chen C, Guo S, Schlegel O, Lutterman J, Ling K, Zhang F: Dual activities of an X-family DNA polymerase regulate CRISPR-induced insertional mutagenesis across species. NATURE COMMUNICATIONS 2024, 15.

42. Edwards K, Johnstone C, Thompson C: A simple and rapid method for the preparation of plant genomic DNA for PCR analysis. Nucleic Acids Research 1991, 19:1349.

43. Clement K, Rees H, Canver M, Gehrke J, Farouni R, Hsu J, Cole M, Liu D, Joung J, Bauer D, Pinello L: CRISPResso2 provides accurate and rapid genome editing sequence analysis. NATURE BIOTECHNOLOGY 2019, 37:224–226.

44. Edgar R: MUSCLE: multiple sequence alignment with high accuracy and high throughput. NUCLEIC ACIDS RESEARCH 2004, 32:1792–1797.

45. Untergasser A, Cutcutache I, Koressaar T, Ye J, Faircloth BC, Remm M, Rozen SG: Primer3-new capabilities and interfaces. Nucleic Acids Research 2012, 40:12.

46. Aesaert S, Impens L, Coussens G, Van Lerberge E, Vanderhaeghen R, Desmet L, Vanhevel Y, Bossuyt S, Wambua A, Van Lijsebettens M, et al: Optimized Transformation and Gene Editing of the B104 Public Maize Inbred by Improved Tissue Culture and Use of Morphogenic Regulators. FRONTIERS IN PLANT SCIENCE 2022, 13.

47. Anand A, Bass S, Wu E, Wang N, McBride K, Annaluru N, Miller M, Hua M, Jones T: An improved ternary vector system for Agrobacterium-mediated rapid maize transformation. PLANT MOLECULAR BIOLOGY 2018, 97:187–200.

48. Zhang Q, Zhang Y, Lu M, Chai Y, Jiang Y, Zhou Y, Wang X, Chen Q: A Novel Ternary Vector System United with Morphogenic Genes Enhances CRISPR/Cas Delivery in Maize1[OPEN]. PLANT PHYSIOLOGY 2019, 181:1441–1448.

49. Hayta S, Smedley M, Demir S, Blundell R, Hinchliffe A, Atkinson N, Harwood W: An efficient and reproducible Agrobacterium-mediated transformation method for hexaploid wheat (Triticum aestivum L.). PLANT METHODS 2019, 15.

50. Rodrigues SD, Karimi M, Impens L, Van Lerberge E, Coussens G, Aesaert S, Rombaut D, Holtappels D, Ibrahim HMM, Van Montagu M, et al: Efficient CRISPR-mediated base editing in Agrobacterium spp. Proceedings of the National Academy of Sciences of the United States of America 2021, 118.

