## Additional File 11 Supplemental figures for "A deep-learning predictive model for Cas12a activity in wheat"

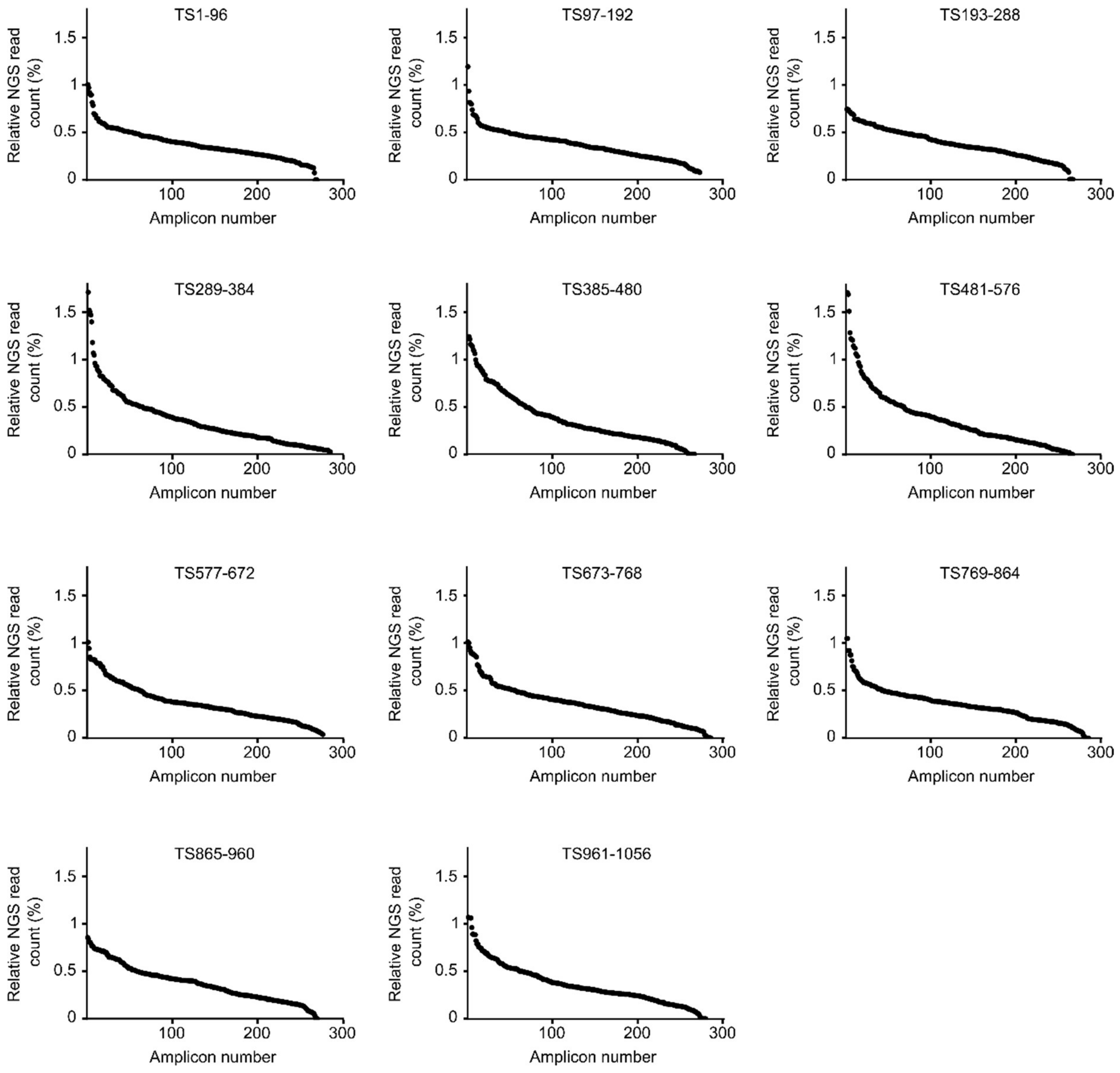


**Fig. S1: Validation primer pair for target site amplicons.**

Validation of genotyping primers by Illumina sequencing and CRISPResso2 analysis. The distribution of the relative abundance of Illumina reads is shown for individual amplicons. Amplicons with <500 reads or >2% background editing were discarded from the screen.

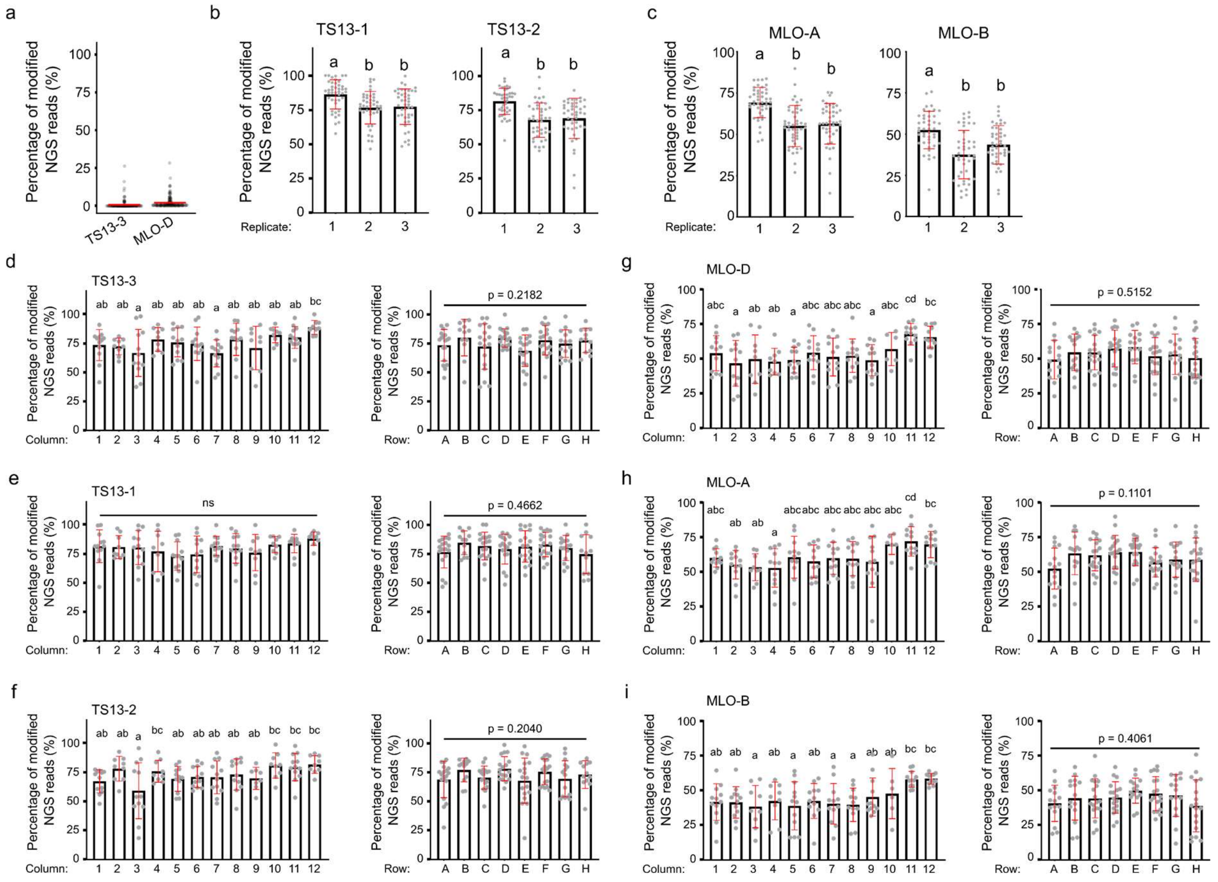

**Fig. S2. Establishment of 96-well RNP transfection and indel frequency variability with transfection replicate and plate position.**

**(a)** Background editing efficiency at TS13-3 and MLO-D. **(b-c)** Influence of the protoplast transfection batch on indel frequency at the off-target sites TS13-1, TS13-2, MLO-A and MLO-B. **(d-i)** Influence of the 96-well plate row and column during protoplast transfection on indel frequency at TS13-3 (d), TS13-1 (e), TS13-2 (f), MLO-D (g), MLO-A (h) and MLO-B (i). Statistics: b-i: One-way ANOVA, Tukey-HSD $p<0.05$.

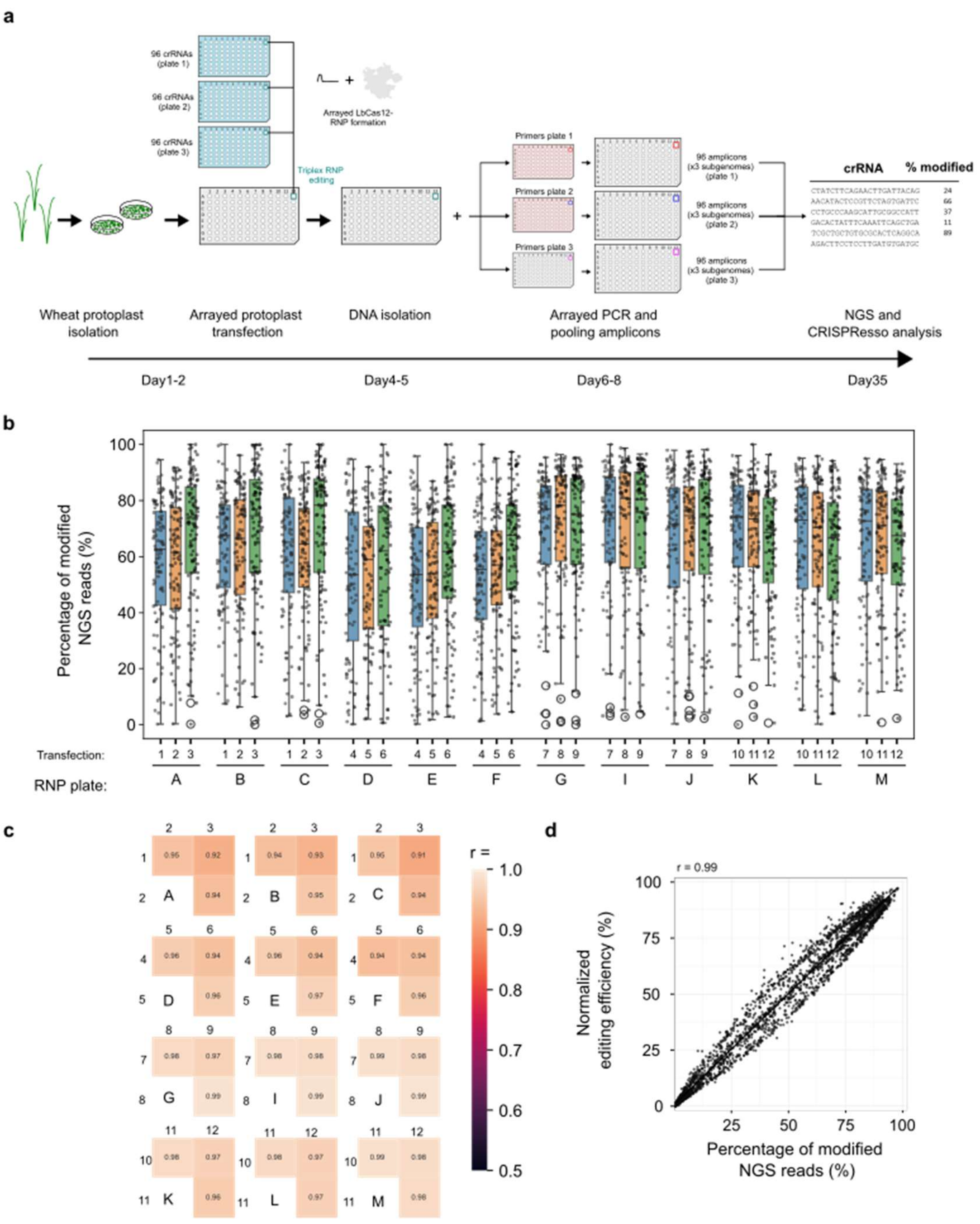


**Fig. S3. CRISPR screen statistics.**

**(a)** Schematic representation of the arrayed CRISPR screen steps and multiplex pipetting scheme. Wheat protoplasts were isolated and three 96-well plates of RNPs were assembled in parallel. Three RNPs were co-transfected per protoplast well and the plate was incubated for 44h. DNA was then isolated from one 96-well plate used as template for three 96-well PCR plate. Individual PCRs amplified simultaneously the three wheat homeologs that were deconvoluted during amplicon-sequencing analysis. **(b)**. Raw editing efficiency of on-targets measured across all plates and biological replicates **(c)** Technical variability as measured by

Pearson correlation (r) observed between the three independent transfections on individual 96-well RNP plates. **(d)** Correlation between ilogit normalized data and raw indel frequencies for the full screen as measured by Pearson correlation.

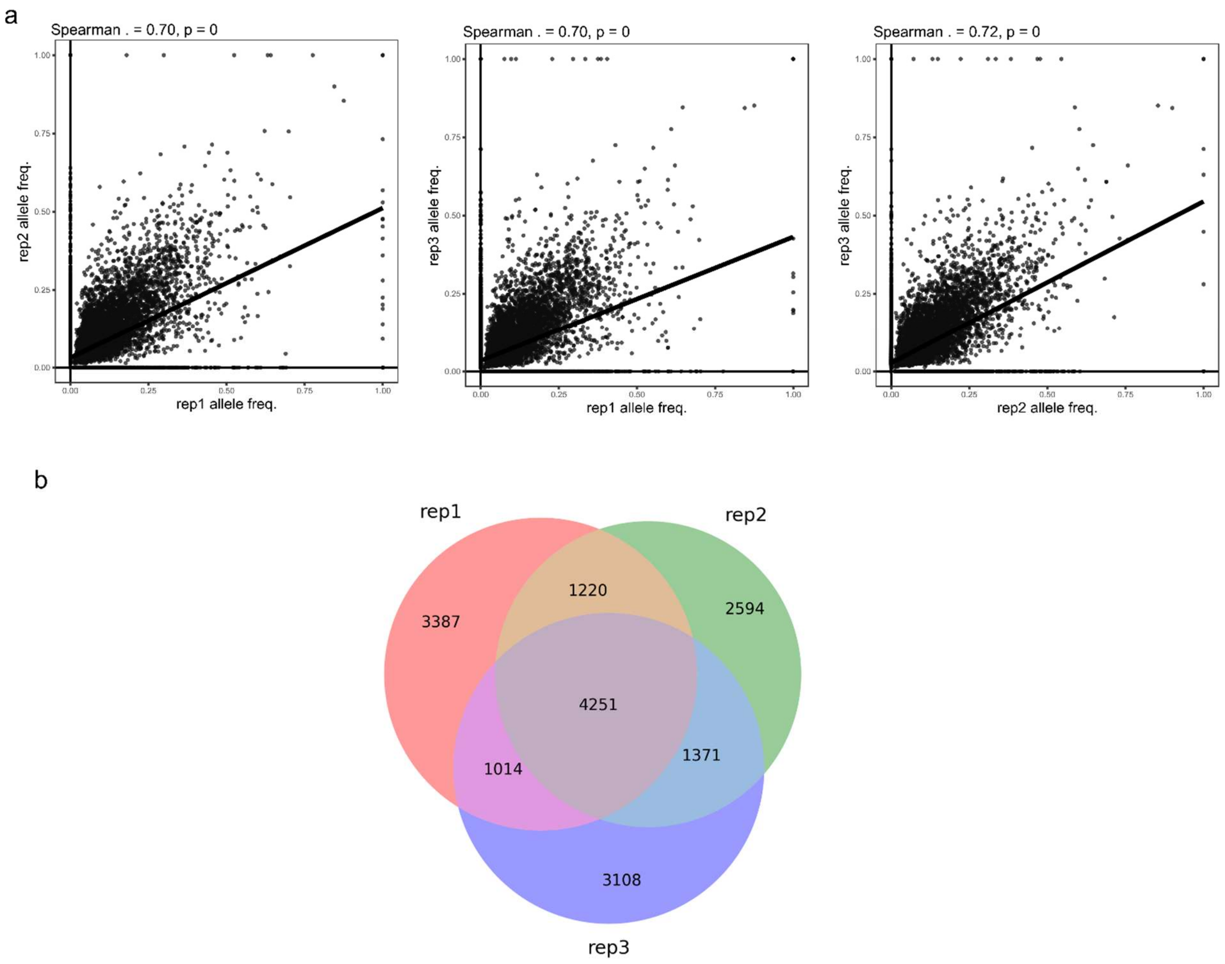


**Fig. S4. Allele overlap between RNP replicates.**

**(a)** Pairwise correlations of relative mutant allele frequencies between the three RNP replicates. All alleles are plotted. Spearman correlations are calculated on alleles that are shared between experimental systems. Samples with a zero value are omitted. **(b)** Venn-diagram showing number of unique mutant alleles shared between the three replicates for all on-targets.

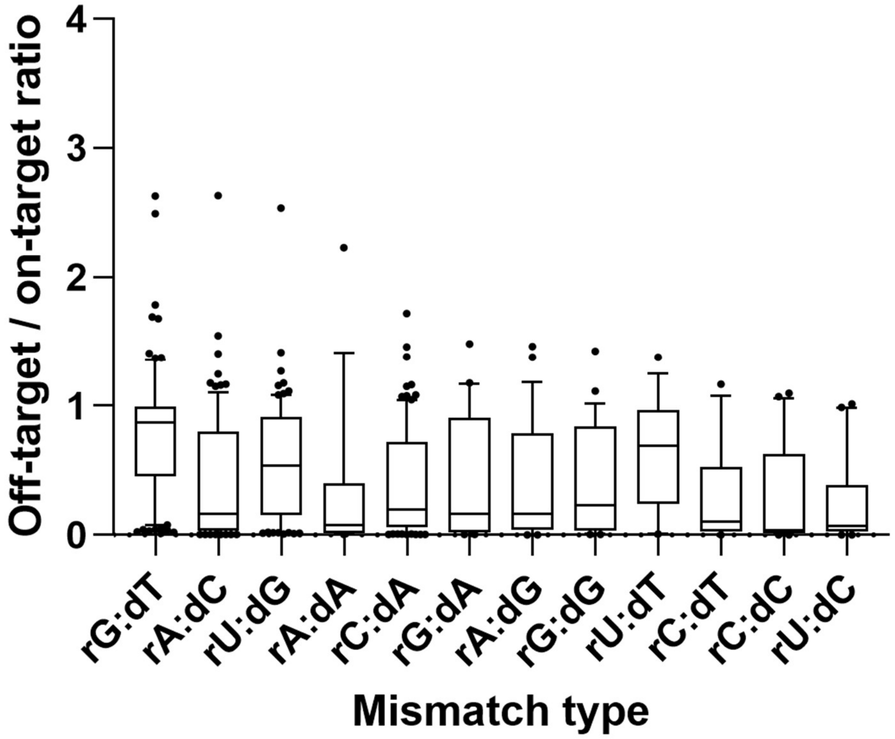


**Fig. S5. RNA:DNA mismatch type affects off-targeting rates.**

Off-target/on-target editing efficiency relative to the type mismatch between the crRNA (r) and DNA target sequence (d). The boxes represent the 25th – 75th percentile, whiskers are 5th – 95th, line is median.

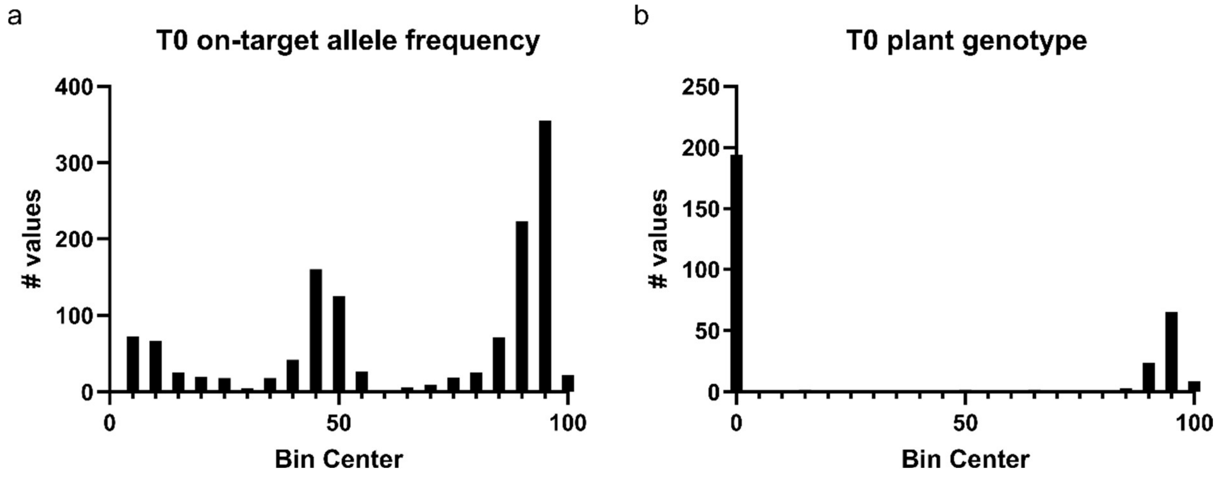


**Fig. S6. T0 plant allele frequencies distribution.**

**(a)** Histogram of the allele frequency of all on-target values for all T0 plants. Values around 0 indicate wild-type, 50 are heterozygous and 100 homozygous. **(b)** Histogram of the on-target genotypes per plant equal to a sum of the allele frequencies in (a). Alleles at 100 indicate both genomic copies are edited, 0 indicated only wild-type alleles.

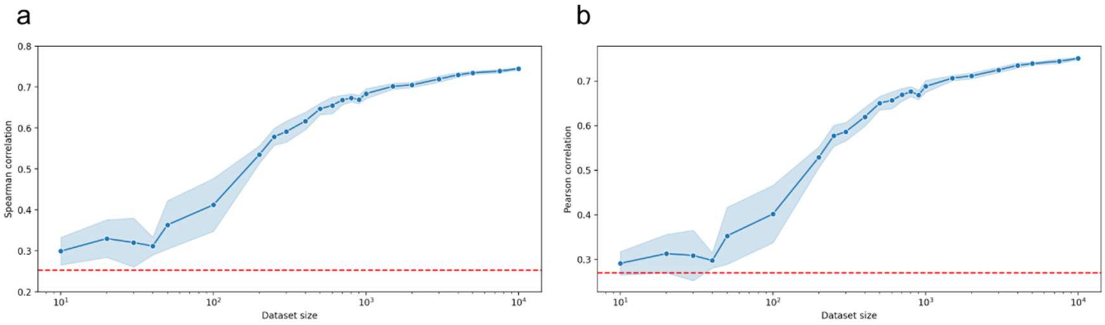


**Fig. S7. 2CaP-5 learning curve.**

**(a-b)** Learning curve analysis for the 2CaP-5 model. Spearman **(a)** or Pearson **(b)** correlation coefficients are depicted in relation to the data set size.
